# Wound-Induced Syncytia Outpace Mononucleate Neighbors during *Drosophila* Wound Repair

**DOI:** 10.1101/2023.06.25.546442

**Authors:** James S. White, Junmin Hua, Jasmine J. Su, Kaden J. Tro, Elizabeth M. Ruark, M. Shane Hutson, Andrea Page-McCaw

## Abstract

In response to injury, cells proliferate, migrate and invade to replace missing cells and close wounds. However, the role of other wound-induced cell behaviors is not understood, including the formation of syncytia (multinucleated cells). Here, we use *in vivo* live imaging to analyze wound-induced syncytia in mitotically competent *Drosophila* pupae. We find that almost half the epithelial cells near a wound fuse to form large syncytia. When the autophagy gene *Atg1* is knocked down, fewer syncytia form, and wounds close more slowly. Further, a computational model of tissue fluidity indicates that cell fusion speeds wound closure time by about one third. Syncytia use several routes to speed wound repair: they outpace diploid cells at the wound margin to lead the initial resealing of the wound; they reduce the need for intercalation as the tissue reshapes during closure; and they pool resources of their component cells to concentrate them toward the wound margin. In addition to wound healing, these properties of syncytia are likely to contribute to their roles in development and pathology.

## INTRODUCTION

Injury is a constant reality of life, and survival requires all organisms to repair wounds. Wound-induced cell behaviors like proliferation, migration, and invasion replace missing cells and close wounds (Martin et al., 2024). Other cell behaviors are induced around wounds, but their contribution to wound healing is not well understood, *e.g.*, the fusion of cells into syncytia. Syncytia are one type of polyploid cell, and it is generally appreciated that increases in ploidy – number of genomes per cell – is a common response of post-mitotic tissues and cells to injury (Losick et al., 2013; Nandakumar et al., 2020; Tamori and Deng, 2013). Wound-induced epithelial syncytia were first observed around epidermal puncture wounds in *Drosophila* larvae and adults (Galko and Krasnow, 2004; Losick et al., 2013), consistent with the idea of polyploidy induction in non-proliferative cells. However, recent studies have observed syncytia around wounds in mitotically competent tissues: around laser-ablation wounds in *Drosophila* pupal epidermis (Wang et al., 2015) and in zebrafish epicardium damaged by endotoxin, microdissection, or laser ablation (Cao et al., 2017). Further, injury associated with the surgical implantation of biomaterials can cause immune cells to fuse into multinucleated giant cells, which are associated with rejection (Al-Maawi et al., 2017). Similarly, injury induces bone marrow-derived cells to fuse with various somatic cells to promote repair (Alvarez-Dolado et al., 2003; Corbel et al., 2003; Davies et al., 2009; Nygren et al., 2004), and injury induces fusions of some *C. elegans* neurons (Ghosh-Roy et al., 2010; Oren-Suissa et al., 2017). The many instances of syncytia being induced by wounds raise the possibility that syncytia offer an adaptive benefit. It is not clear, however, what that benefit is.

Syncytia can form either by endomitosis – mitosis without cytokinesis – or by cell-cell fusion. Cell fusion is widespread during animal development in both vertebrates and invertebrates. For example, myoblasts fuse to form muscle in both vertebrates and in *Drosophila* (Kim et al., 2015a; Lehka and Rędowicz, 2020), epithelial cells fuse during the development of several *C. elegans* tissues including the hypodermis (Podbilewicz and White, 1994) and vulva (Sharma-Kishore et al., 1999), and fusion gives rise to vertebrate syncytiotrophoblasts (Renaud and Jeyarajah, 2022) and multinucleated osteoclast (Søe, 2020). Cell fusions are also observed in disease: pathogen-induced epithelial fusion allows spreading of many viruses including human respiratory syncytial virus and SARS-CoV-2 (Leroy et al., 2020); and the fusion of cancer cells with bone-marrow derived cells is implicated in metastasis (Pawelek and Chakraborty, 2008).

Here, we use live imaging and clonal analysis to understand the behavior of syncytia following laser wounding in the *Drosophila* pupal notum. The unwounded notum is a monolayer epithelium composed of mononuclear diploid cells that are mitotically competent. Nonetheless, during the first several hours after wounding, many of the surrounding cells fuse to form giant syncytia. We previously reported that after wounding the pupal notum, some wound-adjacent cells undergo a single round of endoreplication, increasing their ploidy two-fold (White et al., 2024), but fusion increases ploidy far more, with some wound-induced syncytia observed to have more than a dozen nuclei. Some cell fusions are obvious with apical borders breaking down between cells, while others surprisingly appear as shrinking of a cell’s apical surface. Combined, fusion is a common fate of cells near wounds: about half the cells fuse to form syncytia within 70 µm of a wound with 30 µm radius. To understand the role of fusion-mediated syncytia in wound repair, we undertook two types of analysis, first analyzing an autophagy mutant that reduces fusion after wounding, and second comparing the behaviors of syncytial and mononuclear cells within the same wound. Compared to their smaller mononuclear neighbors, syncytia outpace mononuclear cells to the leading edge. We propose that syncytia accelerate wound repair because cell fusion limits the need to negotiate cell intercalations as the wound closes, a hypothesis supported by computational modeling; and because clonal analysis demonstrates that syncytia mobilize and transport cell resources such as actin from distal cells to the wound margin.

## RESULTS

### A mitotic tissue undergoes cell-cell fusions after wounding

The pupal notum is a monolayer columnar epithelium that will become the epidermis of the adult fly. At the time of our analysis, 12-18h after puparium formation (APF), it sits atop a basement membrane that is still developing (Mehaffey et al., 2024), and it is composed of diploid cells that undergo regular mitotic cycles (Guirao et al., 2015) (Figure 1—figure supplement 1A-C). To analyze cell behaviors around wounds, we live-imaged after laser ablation. Epithelial cell borders were labeled by the adherens junction protein p120ctn-RFP (Ogura et al., 2018) and nuclei were labeled with GFP-labeled Histone2Av (Histone-GFP). Two hours after wounding, we observed what appeared to be prominent syncytial (multi-nucleated) cells around the wound (Figure 1A, Video 1); experiments described below demonstrated that these were indeed syncytia. Some appeared to contain over a dozen nuclei within epithelial borders. For both syncytial and mononuclear cells, it was difficult to assign nuclei precisely to cell borders because notum epithelial cells are not oriented at right angles with respect to the surface, and in 2-D projections, a nucleus was frequently observed outside the cell’s apical border (Figure 1—figure supplement 1D). Accordingly, using apical area and nuclear density, we estimated the number of nuclei within the 3 largest syncytia in different wounds. The number of nuclei in these syncytia increased over time: at 1 h after wounding the three largest syncytia contained an average 3-13 nuclei, and 2 h after wounding they nearly doubled to 6-20 nuclei (Figure 1B). Interestingly, larger wounds generated syncytia with larger apical areas and more nuclei, proportional to wound size, suggesting syncytium formation is a dynamic and scalable response to injury (Figure 1 C-E).

**Figure 1.**
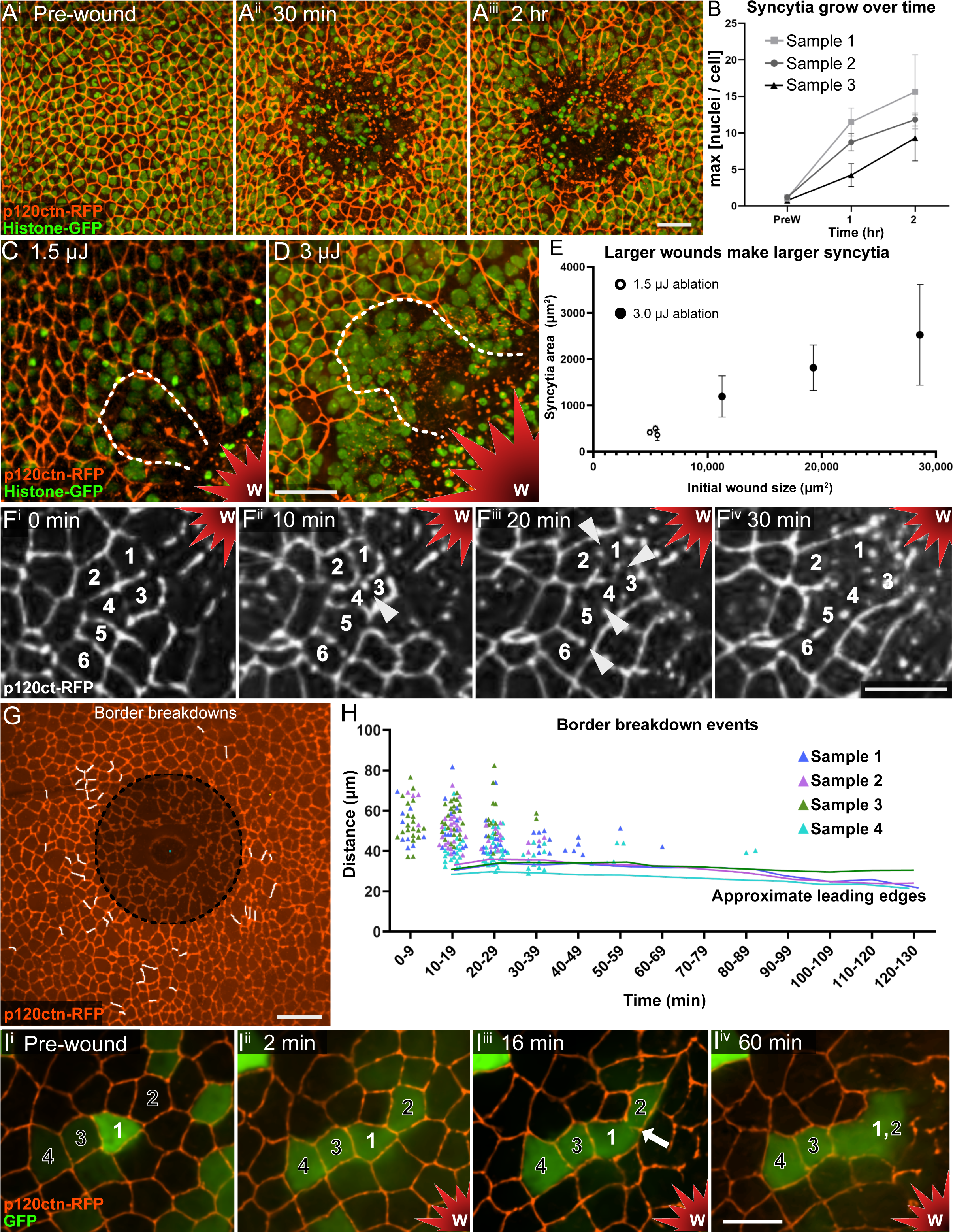
Wounds induce epithelial syncytia via cell fusion. **A)** Syncytia form within 2 h post wounding, evident by the clustering of multiple nuclei within cell borders. **B)** The number of nuclei per syncytia increases over time after wounding. Number of nuclei was estimated based on area and nuclear density (see text) for the 3 largest syncytia of 3 different wounds, mean and SD. **C-D)** Larger wounds generate larger syncytia. Images are at 3 h post wounding, single syncytium outlined. **E)** Syncytial apical area is proportional to initial wound size. Each dot represents a wound, made with either low or high laser energy as sown. For each wound, the mean area of the three largest syncytia is plotted. Bars represent SD. **F)** A time course of six cells fusing within 30 min after wounding. Apical borders are lost (white arrowhead) as syncytia form. Original cells are numbered. **G)** All borders lost to cell fusion (white) mapped to cells in the first frame after wounding. The leading edge of wound closure will form at dashed line; cells within the shaded area were damaged by the wound and will be dismantled. **H)** Distance from the wound center vs time for all border breakdowns in four wounds. Each symbol represents a cell border. Leading-edge locations indicated by solid lines. **I)** Cytoplasmic GFP is expressed in cell 1 before wounding and mixes with neighboring cells 2-4 by 2 min after wounding. Cytoplasmic sharing is followed by the lagging fusion indicator of visible border breakdown (white arrow). The fates of cells 3 and 4 are shown at later times in Figure 2D. Maximum intensity projections in A, C-D, G; single Z slices in F, I. Scale bars: A,C,D,G = 20 µm, F,I = 10 µm. W and red star indicate wound.

Live imaging of cells after wounding revealed the gradual loss of p120ctn along the border between two epithelial cells followed by rounding of the resulting syncytium (Figure 1F,I, and Video 2), such that the initial two-cell morphology gradually became a one-cell morphology. We termed the visible loss of p120ctn “border breakdown”. We determined that as p120ctn was lost, the adherens junction protein E-cadherin was also lost (Figure 1—figure supplement 1E-H) indicating adherens junctions disassembly along the border between the cells. Border breakdowns were found spatially clustered in the first three to four rows of cells, 30-50 μm from the wound center, and they occurred primarily within the first hour after wounding, some within 10 minutes after wounding (Figure 1G,H). These events could be explained by wound-induced cell fusion or by wound-induced epithelial-to-mesenchymal transition. To ask whether border breakdowns represented cell fusions, we analyzed cytoplasmic mixing. Individual GFP-labeled cells (clones) were generated at random locations using the flip-out Gal4 technique (Pignoni and Zipursky, 1997). Before wounding, the level of cytoplasmic GFP fluorescence was stable yet exhibited cell-to-cell variability, allowing some differentiation of cells by intensity (Figure 1I^i^). Minutes after a nearby laser ablation, GFP was observed to diffuse from labeled cells into neighboring unlabeled cells. GFP mixing between two cells was followed by the eventual loss of their shared p120ctn-labeled cell border, confirming that border breakdown is indeed cell fusion, but that cytoplasmic mixing occurs more than 10 minutes before the border breakdown is first observed to start (Figure 1I, Video 2). GFP mixing preceded border breakdowns in all observed cases (n=11). Cytoplasmic GFP mixing resulted in a consistent level of fluorescence across the two cells, indicating that GFP mixing is a reliable indicator of plasma membrane fusion. The order of events demonstrates that plasma membrane fusion occurs many minutes before the visible disassembly of the apically located adherens junctions. Thus, epithelial fusion is a rapid local response to wounding, and border breakdown is a lagging indicator of cell fusion.

### Apical cell shrinking also represents cell-cell fusion, evident later during wound closure

Border breakdowns, representing cell fusion events, occurred mostly within 1 h after wounding, so it was unclear how syncytia grew in size and nuclear number between 1-2 h after wounding. However, an unexpected cell behavior was frequently observed during this time: the apical area of diploid epithelial cells shrank until they disappeared from the epithelial sheet, a phenomenon we termed “apical shrinking” (Figure 2A). Although the appearance of apical shrinking suggested cell extrusion, we were surprised at its high frequency around wounds (Figure 2E), as wounds have already lost many cells and would be expected to prioritize cell survival to speed barrier repair. To understand the fate of apical-shrinking cells, we tracked their nuclei and determined that they did not extrude; rather, all apical-shrinking cell nuclei moved laterally within the plane of the epithelium to join neighboring syncytia (n=7, Figure 2B), suggesting that apical shrinking is an indicator of cell fusion. To better understand this behavior, we analyzed individual GFP-labeled cells so that we could observe the fate of the cytoplasm. Like with border breakdowns, GFP mixing in the X-Y plane of the epithelium was observed to precede the initiation of apical shrinking, although the interval between GFP mixing and apical shrinking was longer, one or more hours (Figure 2C, Video 3). In X-Z projections, GFP labeled cytoplasm associated with apical shrinking shifted laterally to join with a neighboring cell as shown for two apical-shrinking cells in Figure 2D. For the cell labeled with the yellow arrow, before apical shrinking at 60 min post wound, its GFP-labeled cytoplasm extended from the plane of the adherens junctions (Z_AJ_) basally over 8 µm; when apical shrinking was underway at 120 min post wound, its cytoplasm extended less than 5 µm down from the apical plane; and when shrinking was almost complete at 150 min post wound, it extended only 1-2 µm down from the apical plane, appearing as a whisp of cytoplasm connecting the shrinking apical surface with the neighboring syncytia. A similar loss of basal depth over time is observed in the cell labeled with the white arrow. In both these cells, as the cell lost its basal volume, cytoplasm moved laterally to join the neighboring syncytia. Thus, analysis of both nuclei and cytoplasm demonstrated that apical shrinking represented cell fusion events. The spatial distribution of apical shrinking around wounds was similar to border breakdowns, but apical shrinking events occurred later and were more numerous (Figure 2E,F). Thus, cell fusion is evident as either border breakdown or apical shrinking, which both occur after GFP sharing.

**Figure 2:**
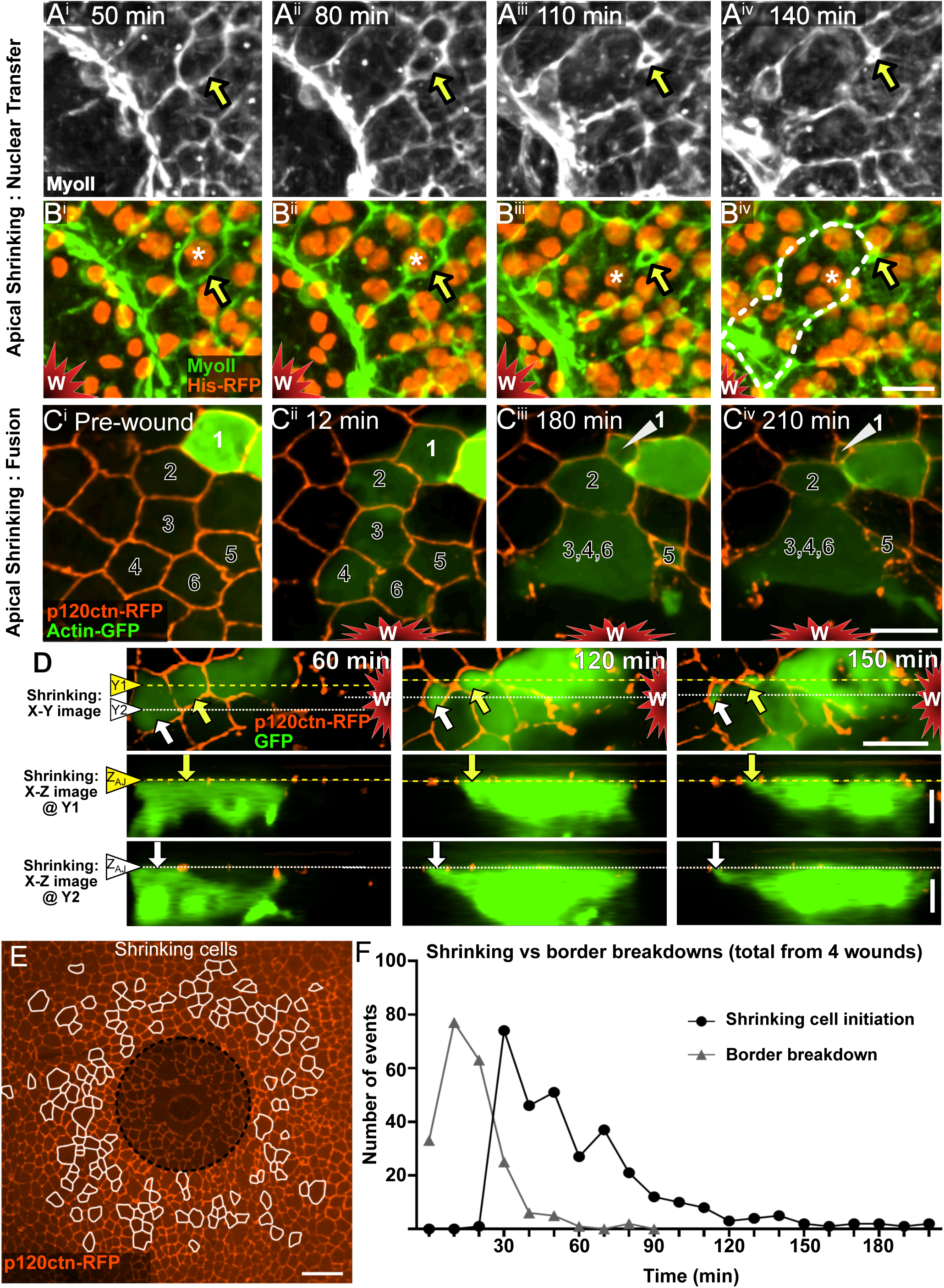
Cell fusion often appears as apical shrinking. **A)** The apical footprint of an epithelial cell shrinks in the epithelial plane after wounding (yellow arrow). **B)** Same sample as A, showing that the nucleus (asterisk) of the apically shrinking cell enters a neighboring syncytium, outlined in B^iv^. **C)** Cell 1 expresses actin-GFP before wounding. After wounding, cell 1 fuses with cells 2-6 evidenced by actin-GFP sharing, then the apical surface of cell 1 shrinks. **D)** Two cells undergoing apical shrinking are identified in the X-Y plane with white and yellow arrows, shown at three times after wounding. Below, two X-Z projections are shown for each, at Y1 and Y2 as indicated, with the Z-plane of the adherens junctions (Z_AJ_) indicated by dotted lines. Rather than extrude, cytoplasm moves to the right as the cell depth in the Z-axis diminishes, so that cytoplasm joins with the neighboring wound-proximal syncytium. These video frames are a continuation of the sample shown in Figure 1I. **E)** All cells displaying apical shrinking mapped to first frame after wounding. **F)** All apical shrinking and border breakdowns were tracked in 3 wounds. Border-breakdown fusions happen sooner after wounding than apical-shrinking fusions. A, B, C, D (top), and E show maximum intensity projections. Lower panels in D show X-Z projections. Scale bar for all panels A and B shown in B^iv.^ A, B, C, D (top),E = 10 µm; D (X-Z projections) = 5 µm.

To determine what percentage of cells around a wound will fuse, we analyzed over 100 single epithelial cells randomly-labeled with cytoplasmic GFP within the radius of observed fusion (80 µm), tracking them for 6.5 h to assess their fate (Figure 3A,B): a full quarter of the cells fused (25%), sharing GFP before borders broke down or apical shrinking; 67% persisted as diploid cells, most with stable GFP, but infrequently (n=3) with GFP mixing and no subsequent cell fusion, suggesting that fusion was initiated then aborted; the remaining 7% could not be tracked (Figure 3—figure supplement 1A). All GFP-labeled cells with apical shrinking had previously shared cytoplasm, indicating that all apical shrinking represents fusion. For the three diploid cells with previous cytoplasmic mixing, we interpret those as cells that initiated fusion but were unable to productively stabilize and expand the site of fusion and instead returned to the diploid state. Fusing cells were strongly skewed toward the center of the wound: within 70 µm, about half the cells (47%) underwent fusion (Figure 3A, left), and fusion continued for over 300 min after wounding (Figure 3—figure supplement 1B,C). Apical shrinking accounts for more than half of all fusing cells (Figure 3C), and the spatial distribution of fusing cells that shrank vs. lost borders was similar (compare Figures 1G and 2E). However, apical shrinking began later than border breakdowns (Figure 2F, Figure 3—figure supplement 1C), continued for several hours after wounding (Figure 2F, Figure 3—figure supplement 1C), and took longer to complete (Figure 3—figure supplement 1D). Thus, fusion associated with apical shrinking accounted for continuing syncytial growth after border breakdowns subsided. Further, cell fusion is a persistent behavior over the course of wound closure.

**Figure 3:**
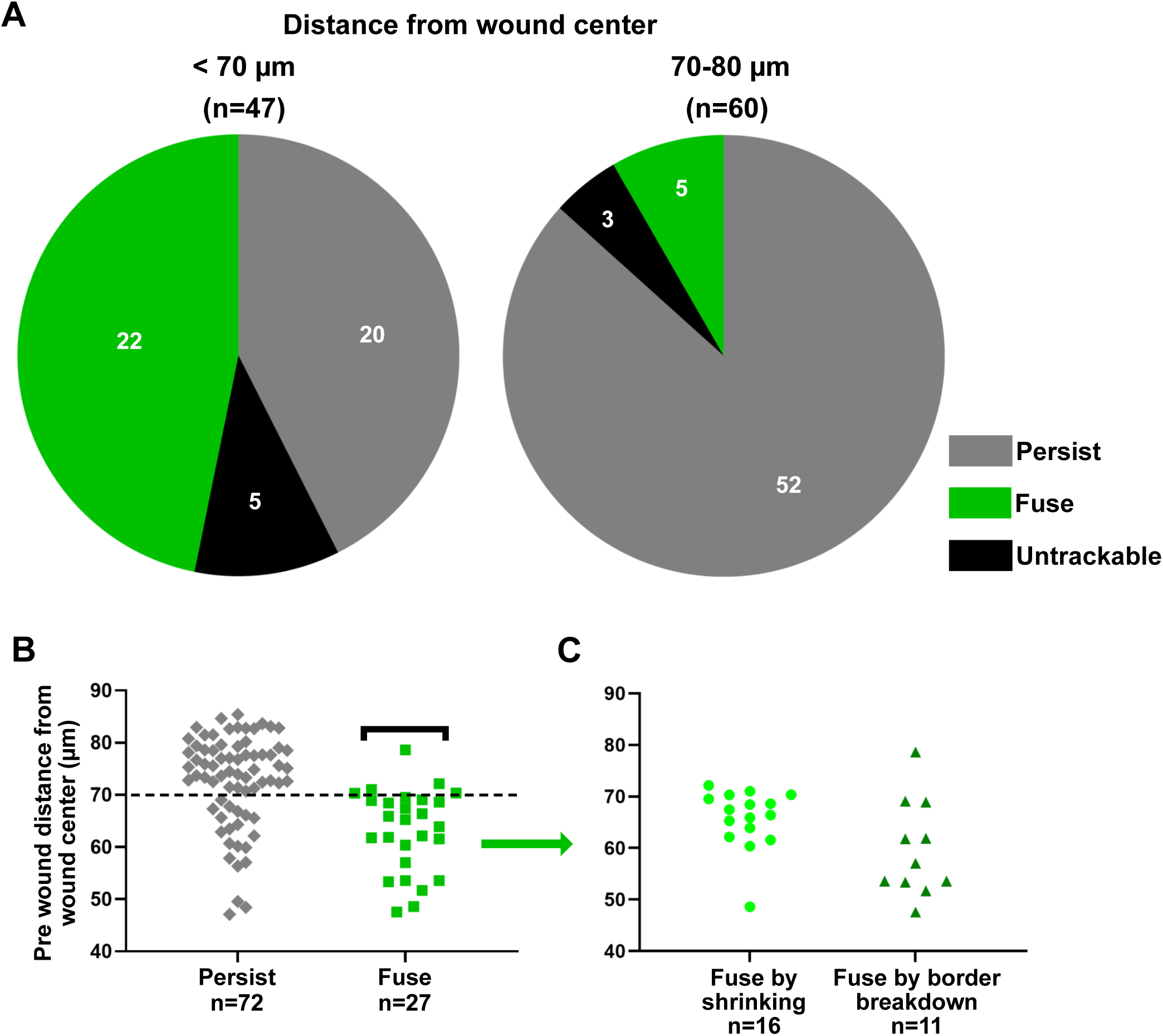
Half the cells near the wound fuse to form syncytia, demonstrated by tracking individual cell fates. **A)** All GFP labeled cells in the region of fusion (80 µm) were tracked in 5 wounds over 6.5 h after wounding to determine frequency of fusion initiation (GFP mixing) and later morphological fusion, *i.e.* a change in the adherens junction marker p120ctn-RFP, which outlined two cells before fusion and one large cell after fusion. All morphological fusion events were preceded by GFP sharing, and 90% of GFP sharing events were followed by morphological fusion. Untrackable cells lost GFP, see Figure 3—figure supplement 1A. **B-C)** Pre-wound location of persisting and fusing cells from panel A is shown with respect to the wound center. Panel B shows that fusion is common within 70 µm. Panel C shows that apical-shrinking fusion and border-breakdown fusion occur at similar distances from the wound.

Our individual cell tracking indicated that syncytia were the result of epithelial cell fusions. However, phagocytic hemocytes are known to migrate to the sites of pupal wounds (Antunes et al., 2013). To determine if hemocytes fused with epithelial syncytia around wounds, we labeled hemocytes with cytoplasmic GFP (*hml>GFP*) and observed their behavior in live imaging. In six wounds, we observed GFP-labeled hemocytes at the basal side of the wound margin; however, we did not observe any events where GFP was transferred from hemocytes to the epithelium, ruling out the fusion of hemocytes with epithelia in pupal wounds (Video 4) .

### Syncytia close wounds faster

To evaluate the role of cell fusion and syncytia in repairing wounds, we wanted to analyze a mutant that blocked syncytia formation. It has been reported that several individual autophagy (*Atg*) genes are required for syncytia formation around small laser ablation wounds in the polyploid *Drosophila* larval epithelium (Kakanj et al., 2022), so we asked if the autophagy gene *Atg1* promotes fusion around laser wounds in the diploid pupal epithelium. Using a dsRNA line known to disrupt syncytia in larvae, we knocked down *Atg1* on one side of the wound expressing *pnr-Gal4*, as illustrated in Figure 4A, reserving the other side of the wound as an internal control, modifying an internally controlled wounding system we have previously reported (O’Connor et al., 2021b). Visualizing Ecad-labeled cell borders across the wound, we found that indeed *Atg1* is required for syncytia formation in pupal wounds, like it is in larval wounds (Figure 4B). Specifically, we observed a reduction in cell fusion, both reductions of border loss and apical shrinking (Figure 4C,D). We next compared wound closure rates on the two sides of the wound (Figure 4E,F). Because leading edge migration is very slightly delayed on the *pnr-Gal4* side of the wound even in control wounds (*pnr>+*, (Hua et al., 2026), we compared the difference in wound radii on the two sides, comparing control wounds to *pnr>Atg1^RNAi^* wounds. By 5h after wounding, the *pnr>+* side lagged behind the other side by only about 10 µm, yet the *pnr>Atg1^RNAi^*side lagged behind the other side by an average of about 25 µm, a reproducible, significant, and large reduction in wound closure when syncytia formation was reduced (Figure 4F). These results suggest that syncytia speed wound closure. It is difficult, however, to disentangle the roles of autophagy and syncytia in this experiment, as the *Atg1* knockdown cells may move more slowly because they are defective in the essential cellular function of autophagy and/or lacking syncytia. To address this question, we found a strong correlation between the size of the cell/syncytium and the distance it had moved from the start, measured at the time when the wound was half-closed. This correlation was true for cells/syncytia in control wounds as well as for those when *Atg1* was knocked down (Figure 4G,H respectively). These data indicate that syncytia formed through cell fusion speed wound closure.

**Figure 4:**
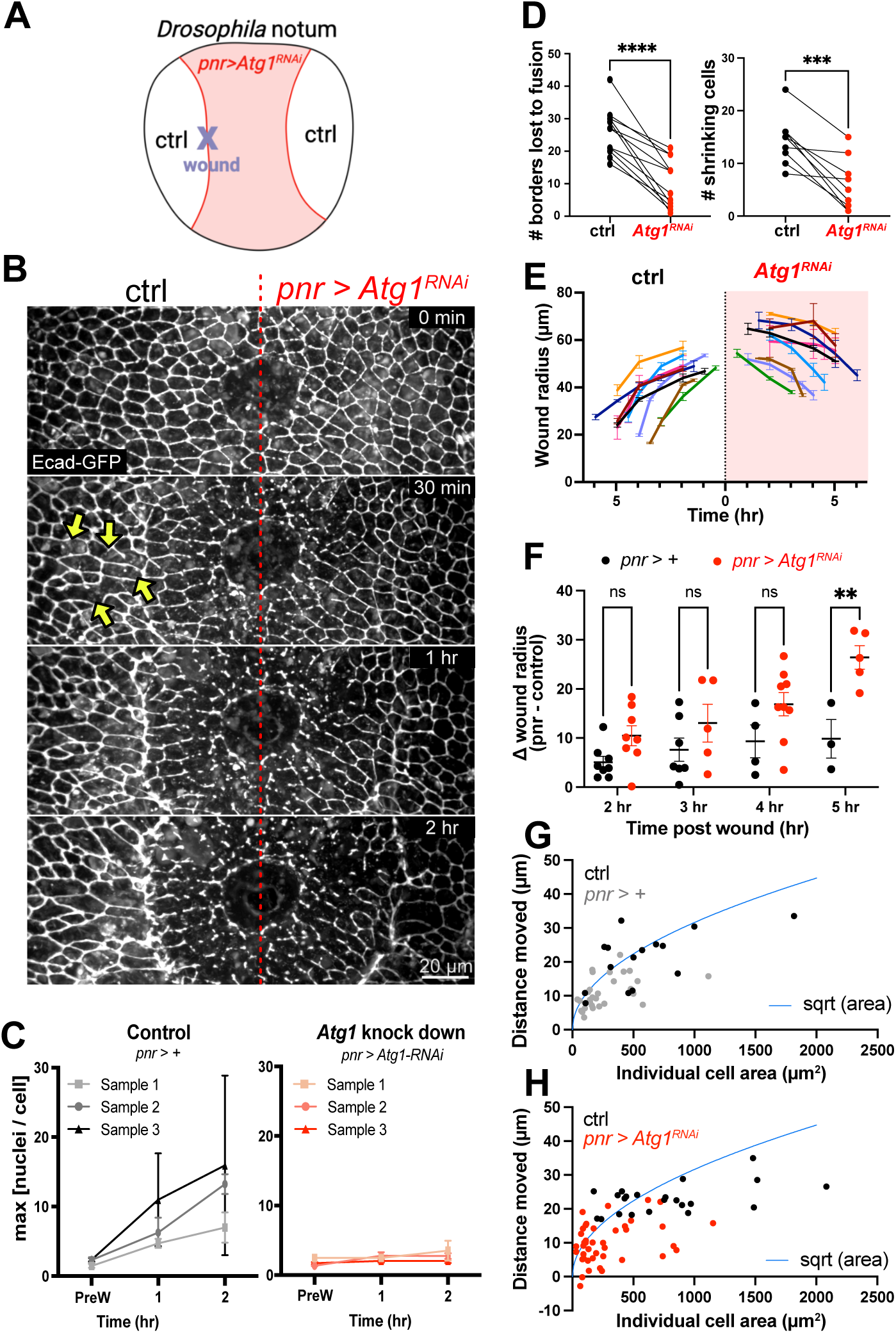
*Atg1* promotes wound-induced fusion and faster wound closure. **A)** Overview: *Atg1* was knocked down in central *pnr* domain (pink), visualized by nls-mCherry (not shown). A laser pulse was targeted to the edge of the *pnr* domain (right or left) so that the wound extended into both *Atg1* knockdown and control domains, allowing the symmetry of response to be analyzed. **B)** Inhibition of *Atg1* decreased the frequency of wound-induced fusion in the *pnr* domain (right) where *Atg1* is knocked down compared to the control domain (left). Some cell fusions are indicated by yellow arrows in the 30 min panel. The red dotted line at the center of each frame indicates the border between the domains, determined by the expression of *UAS-nls-mCherry* (not shown) under control of *pnr-Gal4* on the right. **C)** *Atg1* is required for syncytia to enlarge over time. In the *pnr* domain of control animals, the number of nuclei per syncytia increased over time after wounding. When *Atg1* was knocked down, syncytia remained small over time. The number of nuclei was estimated based on area and nuclear density for the 3 largest syncytia of 3 different wounds, mean and SD. Compare to Fig. 1B. **D)** Inhibition of *Atg1* decreased the number of wound-induced fusions in the *pnr* domain compared to the control domain in the same animal. For border loss, n = 11 pupae, p < 0.0001. For apical shrinking, n=9 pupae, p = 0.0008 (paired T-test). **E-F)** Wounds closed more slowly when *Atg1* was knocked down. (E) Radius of the wound was calculated by the distance between the wound center and the leading edge in *pnr* or control domains. (F) Difference in the wound radius between *pnr* and control domains over time (*i.e.*, wound asymmetry) is shown. Each dot represents one wound. With no genetic manipulation in either domain (black), wound radius was slightly larger in *pnr* than in control domain. When *Atg1* was knocked down in the *pnr* domain, the asymmetry grew over time indicating slower wound closure on the *Atg1* knockdown side. p = 0.0031 (5 hr) calculated from two-way ANOVA, fit full model, comparing each cell mean with the other cell mean in that row. Bar = 1 +/- SEM. **G-H)** Larger cells (*i.e.* syncytia) move further during wound closure, evident in control wounds (G) or with *Atg1* knockdown (H). Each data point represents a cell or syncytium present at the leading edge when the wound is half closed. The curve of square root area was overlaid for comparison, as it represents the amount of movement expected based on reshaping of a cell/syncytium from an initial round state to one strongly elongated towards the wound.

### Syncytia outcompete mononucleate cells at the leading edge of repair

To understand this phenomenon better, we further investigated control wounds. Because about half the cells fused to form syncytia around wounds, we were able to compare the behavior of syncytial to non-syncytial cells within the same wound, providing a well-controlled environment for assessing how syncytia contribute to wound closure. In live imaging, we observed that syncytia frequently moved faster than unfused cells toward the wound, overtaking unfused cells as the wound closed. Figure 5A tracks four GFP-labeled cells near a wound, with cell 1 furthest from and cell 4 closest to the wound (Figure 5A^i^). Middle cells 2 and 3 fuse with five others into a large syncytium, which advances toward the wound faster than both non-fusing cells, and by 400 min after wounding the syncytium extends well beyond the unfused cells (Figure 5A^iii^). This behavior was evident even without GFP labeling: Figure 5B-C show a group of cells at the leading edge of the wound (the wound is recognized by the lack of p120ctn). At 90 min after wounding, this region of the leading edge comprises three mononuclear cells flanked on either side by syncytia (outlined in yellow); the middle of the three mononuclear cells is outlined in orange (5B) and white (5C). By 150 min, the syncytia have pushed out all three mononuclear cells from the leading edge, and at 360 min two of these mononuclear cells are visible about 20 µm back from the wound edge. Indeed, we observed that several hours after wounding, the entire leading edge was occupied by syncytia. To investigate how syncytia came to occupy this position at the front lines of wound healing, we expressed MyoII-GFP/Zip-GFP, which along with actin forms the contractile purse string and makes the leading edge visible, along with p120ctn-RFP to label cell borders. We analyzed the persistence of all unfused and syncytial cells at the leading edge over the course of closure for three wounds, starting when the leading edge was first visible about 30 min after wounding (Figure 5D-G). At the start, 75% of the perimeter was occupied by syncytia with the rest occupied by 10-13 unfused cells (Figure 5D,G). As the wound closed, the syncytia became larger and displaced all the unfused cells, with the last unfused cell removed from the leading edge 20-160 min before closure (Figure 5E-G). No unfused cell persisted at the leading edge through wound closure, indicating that syncytia outcompete unfused cells to close the wound.

**Figure 5:**
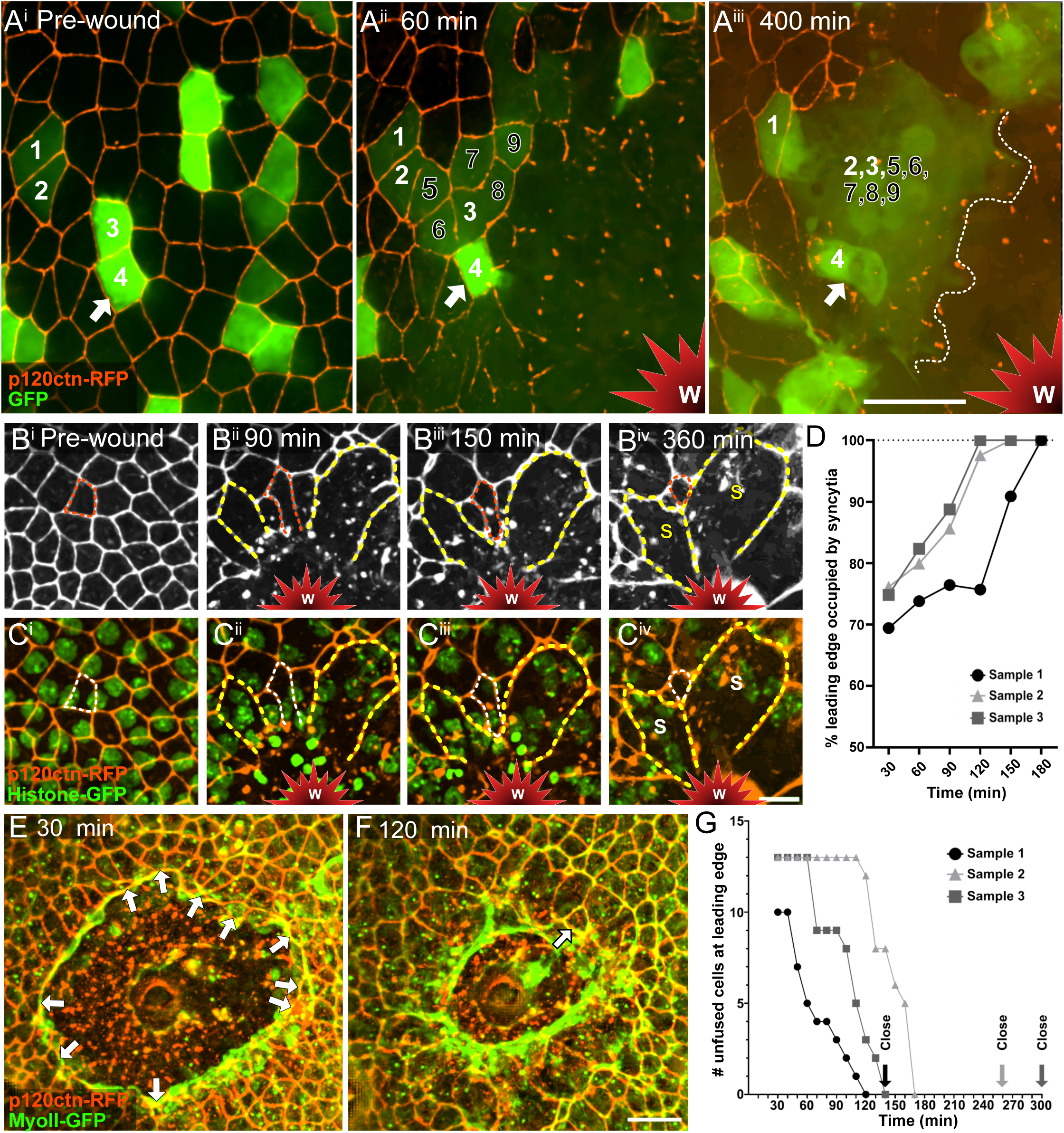
Syncytia outpace mononuclear cells. **A)** Before wounding, two clusters of cells are labeled with GFP, 1-2 and 3-4 (white numbers). After wounding, cells 2 and 3 fuse with neighbors 5-9 to form a syncytium (dashed leading edge in A^iii^), which advances toward the wound, passing unfused cell 4 (arrow). **B-C)** Unfused cell (outlined in orange and white) is replaced at leading edge by neighboring syncytia (S and yellow outline). **D)** Leading-edge perimeter was analyzed over time for three wounds, also analyzed in panel G. Percent leading edge occupied by syncytia increased over time to 100%. **E)** Images of sample 1 from graph D. Unfused cells (arrows) were tracked over the course of wound closure; 30 min after wounding, 10 unfused cells at the wound leading edge are indicated by arrows. **F)** At 120 min, the last unfused cell was ejected from leading edge (arrow). The wound closed at 140 min. **G)** The loss of unfused cells was analyzed over time in the three wounds from panel D. All unfused cells are excluded from leading edge by syncytia well before each wound closed 20-160 min later. Images are single Z slices in A^i^-A^ii^ and maximum intensity projections in A^iii^-F. Scale bar for B,C shown in C^iv^ and for E,F shown in F. Scale bars in A,E,F = 20 µm, B,C = 10 µm.

### Radial border fusion reduces intercalation, the rate-limiting step of wound closure

Why do syncytia move faster? To address this question, we considered the geometry of fusion because different fusion orientations could provide different benefits for wound repair. We observed that fusions sometimes occur between two adjacent cells equidistant from the wound, as diagramed in the top panel of Figure 6A and exemplified in Figure 1I^iii^-I^iv^ (cells 1,2). This breakdown of a radial border produces a syncytium elongated along the wound edge. Alternatively, fusion may occur between adjacent cells at different distances from the wound, as diagramed in the lower panel of Figure 6A and exemplified in Figure 1F (cells 3,4). This breakdown of a tangential border produces a spoke-like syncytium pointing into the wound. To analyze the frequency and timing of these two different axes of fusion, we calculated the orientations of all lost borders in four wounds. (The angle of the lost border can be determined for border breakdowns; it is difficult to determine for apical shrinking because the other fusion partner is hard to identify). In four analyzed wounds, we identified 235 border breakdowns: 39 radial borders and 196 tangential borders (Figure 6B). Thus, there were about five-fold more tangential borders lost to fusion than radial. Nonetheless, the two types had similar temporal and spatial distributions around the wounds (Figure 6—figure supplement 1).

**Figure 6:**
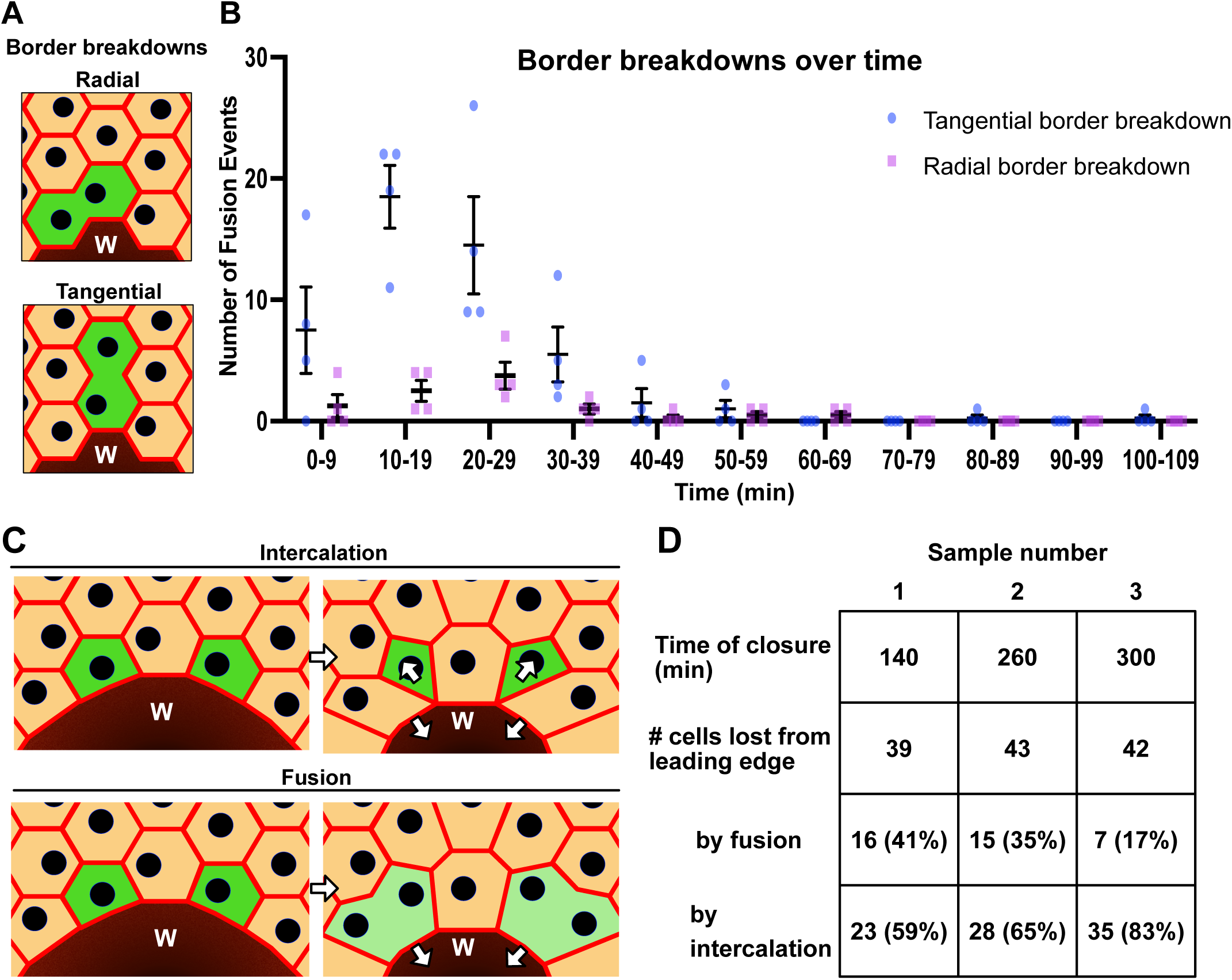
Cells fuse along different axes at different frequencies, reducing the number of wound-proximal cell intercalations. **A)** Illustration of tangential vs radial border breakdown. **B)** More tangential than radial borders break down after wounding. Data are from four wounds, with a total of 235 border breakdowns: 39 radial and 196 tangential. Individual counts are shown per wound with mean and SEM. **C)** Fusions across radial borders reduce intercalation at a wound. **D)** Quantification of intercalations and fusions around the three wounds of Figure 5D,G.

As a wound closes, fewer cells can occupy the ever-smaller leading edge, necessitating a rearrangement of cell adhesions to allow cell intercalation, as depicted in the top of Figure 6C. Indeed, when cells do not fuse after wounding, as in *Drosophila* wing discs, the impact of intercalations on tissue fluidity is known to be a rate-limiting step of wound repair (Tetley et al., 2019). As shown in the bottom of Figure 6C, the requirement for cell intercalation would be reduced specifically by radial border fusion.

To consider the contribution of radial border fusions to wound closure, we took three approaches. First, we analyzed existing data from the three wounds shown in Figure 5D-G, and asked how many leading-edge cells fused radially vs. intercalated: 17-41% of cells removed from the leading edge were removed through fusion, reducing the burden of intercalation substantially (Figure 6D). Interestingly, the larger the percentage of cells that fused rather than intercalated, the faster the wound closed (Figure 6D).

Second, we estimated tissue fluidity as measured by the cell shape index, *ρ* = *P*⁄√*A*, where *P* is cell perimeter and *A* is cell area. This dimensionless index characterizes the cell jamming transition in epithelia. In idealized vertex models, this transition occurs at ρ = 3.81: for lower values, the epithelium is in a jammed state with energy barriers that limit intercalations, giving rise to a solid-like tissue; for higher values, the energy barriers vanish, yielding an unjammed fluid-like tissue (Bi et al., 2016; Park et al., 2015). In real epithelia, ρ is increased by the fact that cell edges are not necessarily straight (as they are by construction in vertex models), and thus the solid-fluid transition may take place at different values of ρ (Wang et al., 2020), but higher ρ still implies greater tissue fluidity. To measure cell shape index, we segmented images of the pupal notum before and after wounding, extracted perimeter and area data for each cell, and calculated the distributions of cell shape index for both syncytia and non-syncytial cells (Figure 7A-B). The mean cell shape index increased significantly after wounding with syncytia having a higher shape index than nearby post-wound non-syncytial cells (*p* = 0.001) and both having higher shape indices than pre-wound cells (*p* < 10^-6^). The post-wound epithelium is thus substantially more fluid-like than it was before wounding

**Figure 7:**
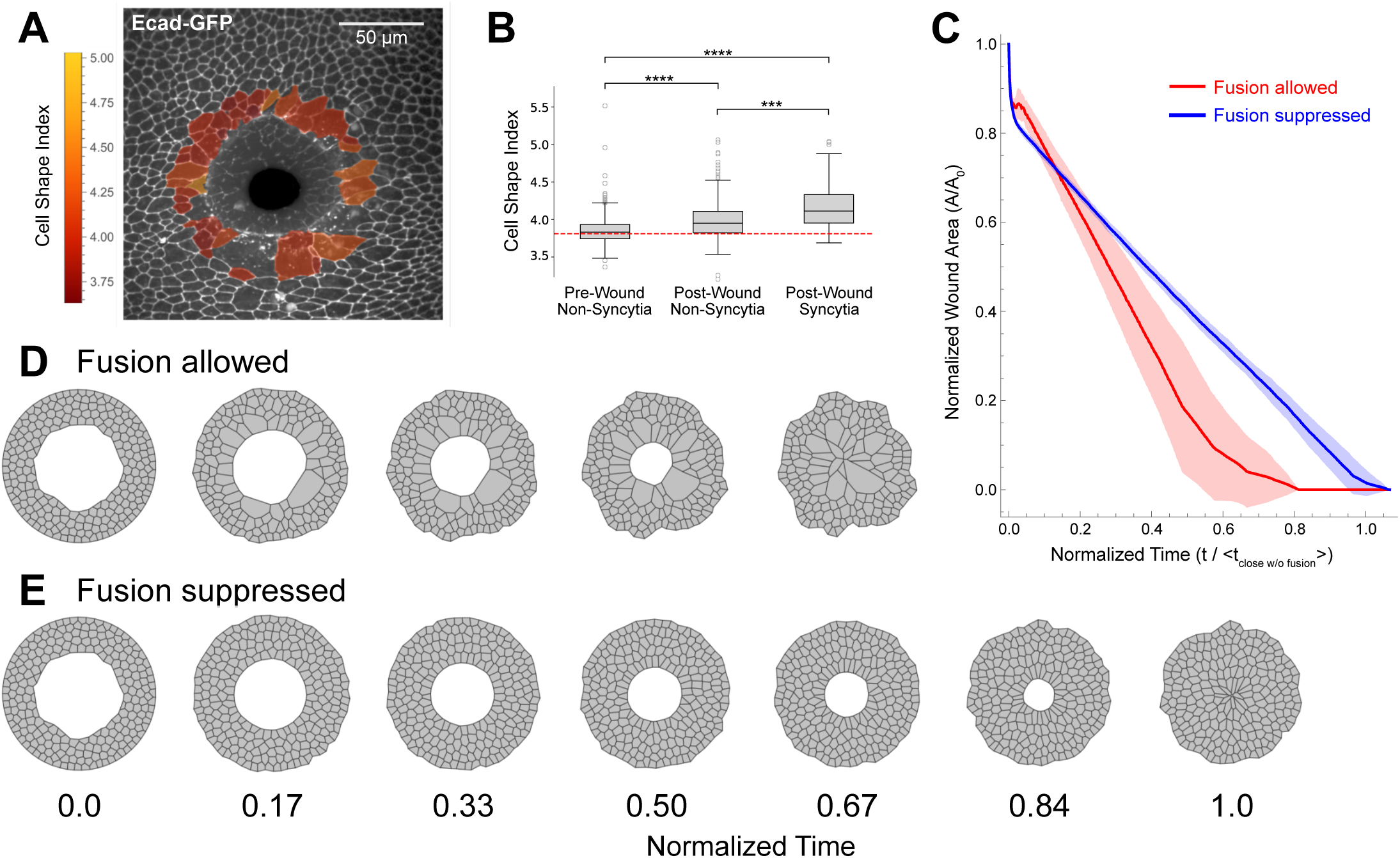
Computational modeling indicates that fusion speeds the rate of wound closure because of reduced cell intercalation. (A-B) Experimental observations of cell shape index (perimeter divided by square root of area). By two hours after wounding, the cell shape index has increased for both syncytia and non-syncytial cells within two rows of the wound margin. These increases indicate an increase in tissue fluidity. (A) Cells and syncytia were segmented, and cell shape index calculated, as shown for this image at 115 min after wounding; cell shape index indicated by color-coding. (B) Pre-and post-wound cell-shape distributions were compiled from 6 pupae. The red dashed line represents the critical cell shape parameter, 3.81, for the solid-to-fluid transition in vertex models of epithelia. Grey boxes span the interquartile range with a line at the median; whiskers extend out 1.5x the median-to-closest-quartile range; and outliers beyond the whiskers are plotted as individual points. Distributions include 1304 pre-wound mononucleate cells, 168 post-wound mononucleate cells, and 52 post-wound syncytia; **** denotes p < 10^-6^ level; *** denotes p < 0.001. **(C-E)** Vertex model simulations of wound closure with cell fusion allowed or suppressed. When the spatial and temporal probabilities for cell fusion match experimental observations, the model yielded 55 ± 22 fusions per wound (mean ± standard deviation), with 15 ± 6 of these occurring at radially aligned edges. As shown in C, fusions in the model speed wound closure: four simulations per condition; shaded regions denote standard deviation; time normalized to the average time to closure when fusion is suppressed. Panels D-E show a series of still frames from a matched pair of simulations in which fusion is allowed or suppressed. Corresponding videos are available as Video 5.

Third, we adapted the computational vertex model from Tetley et al. (2019) to include probabilistic cell-cell fusions to directly investigate the role of cell fusion in tissue fluidity and wound closure. We generated four initial cell configurations with similarly sized wounds and used these to run four matched pairs of simulations – one that proceeded to close without any cell fusions and the other that allowed fusions to occur with spatiotemporal probabilities that matched our experimental observations (Figure 6B, Figure 6—figure supplement 1A). On average, the probabilistic fusion-allowed simulations had 68 ± 14 (mean ± SD) fusions and closed the wound in about 1/3 less time than those without fusions (Figure 7C). Selected frames from one of the matched simulation pairs are shown in Figure 7D-E and full videos of this pair are available as Video 5. Importantly, the modeled syncytia had no special properties: same contractility as other cells, same target shape index, and same target area as the sum of the fused pair. Just the presence of these larger cells reduced the need for intercalations during wound closure and thus sped up the process.

### Tangential border fusions allow resources from distant cells to be mobilized to the wound edge

In contrast to radial border fusions, tangential border fusions might provide a way for cellular resources that would be trapped in distal cells to move toward the wound to contribute to closure. To test this hypothesis, we generated small flip-out clones expressing actin-GFP and wounded such that unlabeled cells intervened between the labeled cell and the leading edge (Figure 8). Like cytoplasmic GFP in Figures 1I and 2C, we observed actin-GFP to equilibrate between cells soon after fusing. For syncytia that did not have access to the leading edge (n =2), actin-GFP levels remained uniform (Figure 8A,B). In contrast, syncytia with access to the leading edge (n = 10) first uniformly distributed actin (Figure 8C^ii^), but once the leading edge was contacted, they redistributed actin-GFP to it (Figure 8C,D). Kymographs of actin-GFP confirm that regardless of location, actin equilibrates between fusing cells within 5 minutes (Figure 8B^iii^, D^iv^); however, nearly all actin that originated in the distal cell is redistributed to the leading edge in a syncytium positioned there (Figure 8D^iv^). In one striking instance, actin-GFP appeared to travel through three cells to arrive at the leading edge from its initial location three rows back (Figure 8E-G, Video 6). Importantly, although actin is labeled from only one of the fusing cells, it likely represents the location of actin from all fusing cells, explaining why syncytia are better able to occupy the leading edge. We envision that other resources would also be concentrated in subcellular regions as needed by syncytia. This ability of syncytia to concentrate pooled actin to the leading edge may allow syncytia to outcompete their smaller mononuclear neighbors and close wounds more rapidly.

**Figure 8:**
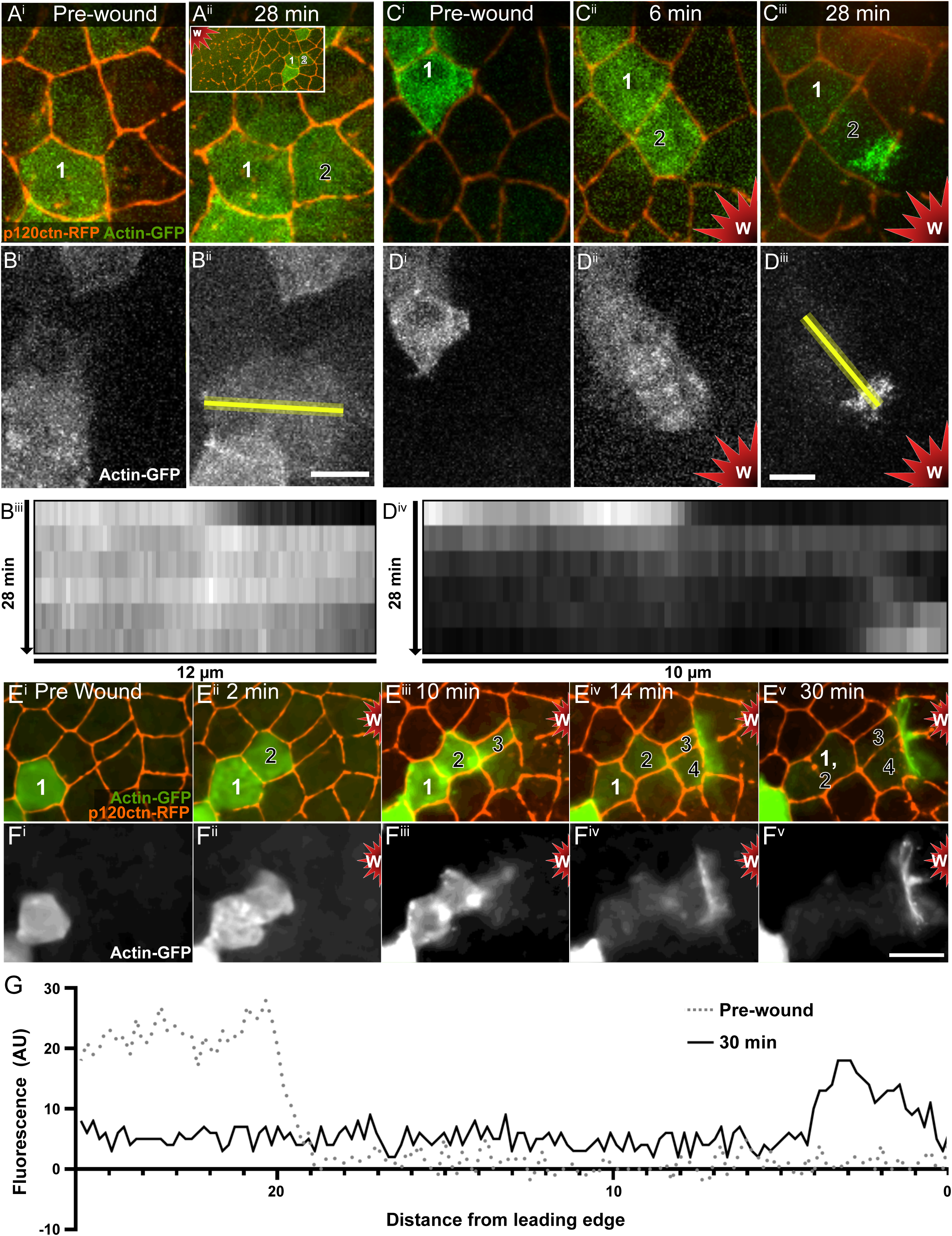
Syncytia concentrate pooled resources at the leading edge. Random scattered cells expressing actin-GFP were generated by heat-shock mediated flip-out expression of Gal4. **A-B)** Labeled actin is expressed in cell 1 before wounding (A^i^,B^i^). These cells are not close to the wound (inset in A^ii^). By 28 min after wounding, actin-GFP equilibrated between cells 1 and 2 (A^ii^,B^ii^), demonstrating cytoplasmic fusion. The resulting syncytium had no access to the leading edge, and actin remained equilibrated, as shown in the kymograph (B^iii^) generated from actin-GFP intensity over time at the yellow line in B^ii^. Fusion was confirmed at 180 min after wounding by apical shrinking of cell 2 (not shown). **C-D)** Labeled actin is expressed in cell 1 before wounding (C^i^,D^i^) and equilibrates between cells 1 and 2 by 6 minutes after wounding, demonstrating cytoplasmic fusion (C^ii^,D^ii^). The resulting syncytium contacts the leading edge, and by 28 min after wounding actin-GFP from cell 1 is redistributed to the wound margin (C^iii^,D^iii^), as shown in the kymograph (D^iv^) of actin intensity over time at the yellow line in D^iii^. Fusion was confirmed at 55 min after wounding, by apical shrinking of cell 1 (not shown). **E-F)** Before wounding, actin-GFP in cell 1 is three cells away from the future leading edge. After wounding, fusion of cells 1-4 allows actin-GFP to be subcellularly localized to the leading-edge actin cable. Border breakdown is visible between cells 1-2 at 30 min after wounding (E^v^) and apical shrinking occurs later (see Video 6). **G)** Mean profile plot of actin-GFP comparing cell 1 before wounding (from panel F^i^, dotted line) to the syncytia of cells 1-4 at 30 min after wounding (from panel F^v^, dark line), demonstrating that nearly all actin-GFP has been relocalized to the leading edge from its starting position 20-30 µm away. Single Z slices shown for E^i-ii^, F^i-ii^; maximum intensity projections for A-D, E^iii-v^, F^iii-v^. Scale bar for panels in A-B shown in B^ii^, 10 µm. Scale bar for panels in C-D shown in D^iii^, 5 µm. Scale bar for panels E-F shown in F^v^, 10 µm.

## DISCUSSION

Previous work established that polyploid cells – both cells with multiple nuclei and cells with enlarged nuclei – are important for closing epithelial wounds in post-mitotic tissues (Losick et al., 2013). Here we have used live-imaging to analyze the response to wounds of a diploid mitotic tissue, the epithelial monolayer of the pupal notum. We found that syncytia form around wounds by cell fusion at a remarkably high rate, with almost half the epithelial cells within 5 cells of the wound fusing with neighbors over the course of hours. Syncytial size increases with wound size, indicating syncytia formation is a dynamic and scalable response to wounding. Several lines of evidence demonstrate that syncytia are faster at closing wounds than unfused cells. Knockdown of the autophagy gene *Atg1* reduces fusion around pupal wounds as was previously reported for larval wounds; *Atg1* knockdown also reduces the rate of wound closure. In both *Atg1* knockdowns and in controls, there was a consistent observation that the bigger the syncytium, the faster it moved. Within control wounds, syncytia completely displace unfused cells of the same genotype at the leading edge of the wound, such that wounds are closed entirely by syncytia, even though there is a smaller number of syncytia than unfused cells. Finally, computational modeling of wound closure with and without cell fusion shows that wounds with fusion-based syncytia close faster because of the reduced need for cell intercalations.

Thus, wound-induced polyploidy is a generalized strategy for wound closure, used by both mitotic and non-mitotic tissues. Although one might have wondered if polyploidy was a wound response limited to tissues that cannot use mitosis to increase nuclear content, both endoreplication and cell fusion into syncytia occur after wounding in this mitotically active diploid tissue (Wang et al., 2015; White et al., 2024; this study). The requirement for *Atg1* for fusion in both polyploid larval tissue and in diploid pupal tissue suggests that the fusion mechanisms are common, despite the altered cell cycle of the starting cells. Further, our finding that larger syncytia travel faster suggests that syncytia – and by extension perhaps cell enlargement via endoreplication –occur as adaptations to close wounds faster.

We identified two unique properties of syncytia that may improve their wound-closing ability. First, syncytia are larger than neighboring diploid cells and their presence enhances tissue fluidity. Previous studies have determined that the rate-limiting step in epithelial wound healing is the process of cell intercalation or rearrangement (Tetley et al., 2019). As the wound closes and the leading edge encircling the wound gets smaller, fewer cells can occupy the leading edge. The cells leaving the leading edge must remodel their cellular junctions, requiring time and energy. Overall, tissue solidity slows closure, so enhancing fluidity promotes faster closure. By fusing into larger syncytia, cells reduce the need for adhesion remodeling and intercalation and instead just change shape as the leading edge becomes smaller. Moreover, the resulting larger cells can have significantly more fluidity and less epithelial tension than their diploid progenitors, as recently demonstrated in syncytia formed by age-induced epidermal cell fusion (Dehn et al., 2023). As our computational modeling demonstrates, fusion can speed wound closure by ∼33% even when the resulting syncytia have no special properties besides being larger than neighboring diploid cells. Making the syncytia individually more fluid would be expected to speed closure even more.

The second unique property of syncytia is their ability to pool and concentrate the cellular resources of the many constituent cells into a subcellular location within the large syncytium. We explored this concept by monitoring the localization of actin within syncytia. Actin is important for wound healing, as epithelial wounds close through the action of two actin-dependent processes: the cinching of a multicellular actin cable at the leading edge that surrounds the wound, sometimes called a purse string (Abreu-Blanco et al., 2012; Bement et al., 1993; Martin and Lewis, 1992; Tamada et al., 2007), and the protrusion of actin-rich projections into the wound (Abreu-Blanco et al., 2012; Farooqui and Fenteany, 2005). Pupal wounds have been shown to use both processes, with the actin cable developing early and actin protrusions appearing later in wound closure (Antunes et al., 2013). By labeling actin in individual cells (clones), we were able to follow it redistribution in syncytia after fusion. As expected, at first labeled actin diffuses and equilibrates throughout the new large cell. Remarkably however, when the syncytium is in contact with the leading edge, labeled actin is concentrated at the leading edge, even if the actin originated several cells away from the leading edge. Without fusion, the labeled actin would have been trapped in a distal cell without access to the leading edge, but after fusion, the syncytium can concentrate the actin of its many component cells at the wound front. If N represents the number of cells that fused, our results suggests that syncytia can apply up to N times more actin to the leading edge; considering that we observed syncytia with dozens of nuclei, this could represent a significant enhancement of actin at the leading edge. Increased actin might explain the ability of syncytia to outcompete diploid cells at the leading edge. Presumably, other resources such as mitochondria and ribosomes could also be pooled and concentrated by syncytia at cellular locations where they promote wound healing. It is known that mitochondrial fragmentation is localized to the site of cellular injury (Horn et al., 2020), and that ribosomes are localized at the tips of severed neurons (Noma et al., 2017), and syncytia would have access to a larger pool of mitochondria and ribosomes than their component cells. By pooling the cellular resources of component cells, syncytia may also allow lethally damaged cells to survive by providing them with needed survival factors originating in cells further from the wound. Thus, the concept of resource sharing that we demonstrate with actin may have ramifications for many resources. Interestingly, it has been demonstrated that wound-induced endoreplicating cells also increase protein levels of some wound-repair resources, such non-muscle myosin, presumably possible because of the many copies present in the polyploid genome (Losick and Duhaime, 2021).

Two visibly distinct cellular behaviors attended fusion in the apical plane of the adherens junctions: border breakdowns and apical shrinking. Both processes were confirmed as fusion by expressing GFP in clones and analyzing GFP sharing at fusion. Border breakdowns occurred sooner after wounding than apical shrinking and appeared to be a faster process, as apical shrinking started later and took hours to complete. It is unclear whether these differences in appearance and timing indicate different mechanisms, as both require the autophagy gene Atg1. For border-breakdown, it is simple to envision that fusion initiates near the apical adherens junctions, which are under more tension than the basolateral membranes, and this may explain why fusion proceeds more quickly there. We recently found that although epithelial tension drops after laser wounding, it is restored within about 10 min (Han et al., 2023), consistent with the timing of apical border breakdown. Also consistent with this model, in the *C. elegans* hypodermis, developmental cell fusion has been observed to initiate at the apical side and travel in the apical-to-basal direction (Mohler et al., 1998). It is also possible that fusion initiates at a more basal site, and that fusion proceeds apically, with adherens junction disassembly occurring later. Regardless of the site of fusion initiation, fusion seems to be a relatively equal process when border breakdown is observed, with the resulting syncytium occupying the same footprint that the two fusion partners did, at least initially. For apical shrinking, however, one cell appears to transfer its cytoplasmic contents and nucleus into another recipient cell, with both cytoplasm and nucleus moving laterally to the location of the fusion recipient, while the apical surface of the donor cell shrinks. It is possible that apical shrinking may indicate a different location for the initial fusion pore below the plane of the adherens junctions; it is also possible that this may reflect a difference in the starting pressure of the two fusion partners, so that one subsumes the other.

It is unclear what triggers either border breakdown or apical shrinking fusion. Many developmentally programmed cell fusions are mediated by fusogens, cell surface proteins that bring opposing membranes into close contact with each other, as in the *C. elegans* hypodermis (Chernomordik and Kozlov, 2008; Iosilevskii and Podbilewicz, 2021; Markvoort and Marrink, 2011; Mohler et al., 2002; Podbilewicz et al., 2006; Sapir et al., 2007; Shemer et al., 2004). For other cell fusions, the fusogen is elusive and may not exist, as none has been identified yet in *Drosophila* myoblast fusions, which occur when a fusion competent myoblast generates actin-rich podosome-like membrane protrusions that invade a founder cell (Kim et al., 2015b; Lee and Chen, 2019; Petrany and Millay, 2019; Rushton et al., 1995; Sens et al., 2010). Wound-induced fusion may be a response to the plasma membrane damage that occurs around wounds; indeed, plasma membrane damage has been documented around both laser wounds and puncture wounds (McNeil and Steinhardt, 2003; Shannon et al., 2017). Interestingly, in the large syncytial cells of the *C. elegans* hypodermis, plasma membrane repair after a puncture wound requires the fusogen EFF-1 (Meng et al., 2020).

Polyploidy as a wound response has begun to get increased recognition. In adult *Drosophila,* epithelial puncture wounds are repaired by both endoreplication and syncytia formation (Bailey et al., 2020; Besen-McNally et al., 2021; Grendler et al., 2019; Losick, 2016; Losick et al., 2013; Losick et al., 2016). In the zebrafish epicardium, genetic ablation is repaired by a wavefront of multinucleated polyploid cells formed by endomitosis, and these lead diploid cells to encompass the heart (Cao et al., 2017). In adult mammalian cardiomyocytes, polyploidy may be an adaptive response to maintain growth after the cardiomyocytes lose their ability to complete mitosis. Mouse cornea endothelial cells endoreplicate to increase polyploidy to restore tissue ploidy following genetic ablation (Losick et al., 2016). Mammalian hepatocytes are known to become increasingly polyploid with age (Carriere, 1969; Wheatley, 2008) and in response to various types of injury and disease (Gentric et al., 2015; Madra et al., 1995; Muramatsu et al., 2000; Sigal et al., 1999; Toyoda et al., 2005; Wilkinson et al., 2019). All mechanisms that promote polyploidy – fusion, endomitosis, endoreplication – result in larger cells with the potential to localize more resources; of these, fusion would act the fastest after wounding because there is no need for DNA replication. Interestingly, in *Drosophila* embryos, wounds induce the surrounding cells to become larger by increasing their volume alone and not their ploidy (Scepanovic et al., 2021), suggesting that simply an increase in size is important. Although many examples of wound-induced polyploidy exist, it is still likely to be an underreported phenomenon, as endpoint analysis might miss a transient polyploid response to injury; live imaging is the surest way to identify a polyploid wounding response.

Another polyploid response to injury can occur after surgical implantation of biomaterials for the purpose of guiding regeneration, as reviewed previously (Al-Maawi et al., 2017). In some cases, only mononuclear cells of the immune system respond to the implant, and in these cases the biomaterial is integrated into the body; in other cases, the implant triggers the fusion of immune cells into multinucleated giant cells, and in these cases the material is degraded and rejected. These multinucleated giant cells seem to share properties with the syncytia of the pupal notum, as they are formed by fusion in response to an environmental trigger and they have an aggressive ability to protect the animal in response to wounding. As these studies highlight, understanding the formation, maintenance, and regulation of polyploid cells may improve our ability to successfully implant biomaterials to aid tissue regeneration.

It is often noted that wound responses are similar to cancer cell behaviors. This similarity extends to wound-induced syncytia and their counterparts, polyploid giant cancer cells, as both cell types are highly aggressive and invasive. Chemotherapeutics induce the formation of polyploid giant cancer cells (Illidge et al., 2000; Mosieniak et al., 2015; Ogden et al., 2015; Wang et al., 2013), and some studies indicate that they can form through cell-cell fusion in tumors (Melzer et al., 2018; Noubissi et al., 2015; Powell et al., 2011; Song et al., 2021; Zhang et al., 2021). Once formed polyploid giant cancer cells are hypothesized to escape further chemotherapy treatments due to increased resistance to genotoxic stress (Weihua et al., 2011). These polyploid giant cancer cells and their progeny also exhibit increased migration and invasion potential (Qu et al., 2013; Zhao et al., 2021). The parallels between the behaviors of polyploid giant cancer cells and the wound-induced syncytia of the pupal notum highlight the importance of understanding wound induced syncytia formation in a highly reproducible system, as a basic understanding of how these cells form in the *Drosophila* notum could inform how they become dysregulated in cancer.

## FIGURE LEGENDS

**Video 1: Syncytial cells form after wounding in the *Drosophila* pupal notum.** Epithelial cell borders in red (p120ctn-RFP) and nuclei in green (Histone-GFP). White box on first frame denotes field of view in Figure 1A. Video begins before wounding and extends to 8 h after wounding.

**Video 2: GFP mixing precedes border breakdown after wounding.** Arrow in first frame points to cell border between cells that will fuse after wounding. GFP diffusion into the unlabeled cell precedes visible border breakdown. w, wound region. Cell borders are labeled with p120ctn-RFP. Video begins before wounding and extends to 2 h 10 min after wounding. Same cells as Figure 1I and 2D.

**Video 3: Shrinking cells contribute to syncytia.** Arrow in first frame points to an individual cell labeled with actin-GFP that fuses by shrinking after wounding. This cell contributes its actin-GFP to neighboring cells minutes after wounding then shrinks much later; shrinking is first evident about 1.5 h after wounding and is nearly complete by 3.5 h after wounding. w, wound region. Cell borders are labeled with p120ctn-RFP. Same cells as Figure 2C.

**Video 4:** Blood cells labeled with *hml-Gal4, UAS-GFP* were observed migrating to and along the basal side of the wounded pupal notum. Video is a Z-stack maximum projection. Wound is indicated by w in the first frame. No GFP was observed to enter the epithelium, indicating that blood cells do not fuse with the wounded epithelium. Six similar wounds were analyzed, with no evidence of fusion.

**Video 5**: Computational vertex models show that incorporating cell fusion speeds wound closure. When fusion occurs with spatiotemporal and edge-orientation probabilities that match experimental observations, wound closure in the model with fusion is 33% faster than when fusion is suppressed.

**Video 6: Syncytia pool actin and concentrate it at the leading edge of repair.** An individual cell labeled with actin-GFP fuses with wound proximal cells and contributes its actin to the leading edge of the syncytium. w, wound region. The original source cell and its immediate neighbor go on to shrink into wound proximal cells. Cell borders are labeled with p120ctn-RFP. Same cells as Figure 6E,F.

## Acknowledgements

We thank James O’Connor for his original observations of syncytial cells around pupal wounds, Kalyssa Platt for assistance in data analysis, Kimi LaFever Hodge for technical assistance, Patrick Page-McCaw for confirming data analysis, and Chloe Hecht and Kimi LaFever Hodge for comments on the manuscript. Stocks obtained from the Bloomington Drosophila Stock Center (NIH P40OD018537) were used in this study. JSW was supported by NIH T32HD007502 to Chris Wright, and JH was supported by American Heart 25PRE1374646 to JH. This work was supported by the National Institute of General Medical Sciences R01GM130130 to APM and MSH.

## Resource availability

### Lead contact

Requests for fly lines, reagents, and additional questions should be directed to Dr. Andrea Page-McCaw

### Materials availability

Fly lines generated in this study are available from the Bloomington *Drosophila* Stock Center or from the lead contact.

### Data availability

All primary microscopy videos are available through OSF.

### Experimental model and subject details

#### Drosophila melanogaster

*Drosophila* lines used in this study are in Table S1. All *Drosophila* lines were maintained on standard cornmeal-molasses media supplemented with dry yeast. All flies, except those used in heat-shock flip clonal analysis, were raised at room temperature. For clonal analysis experiments flies were raised at 18 degrees Celsius until the 3^rd^ instar stage when they were heat shocked in a circulating water bath at 37°C for 3 minutes. They were then allowed to develop at room temperature to 15-18hr after puparium formation (APF) before wounding experiments (described below) were conducted. To knock down *Atg1*, *Atg1^RNAi^*was crossed to the stock for internally controlled wounding analysis, expressing *Ecad-GFP* and *TubP-Gal80^ts^* ubiquitously and *pnr-Gal4, UAS-nls-mCherry* in the central region of the notum, similar to our previous system (O’Connor et al., 2021b). Crosses were moved to 29° C for 6 days to allow expression of the RNAi construct and dissected between 12-15 hr APF.

### Method details

#### Pupal mounting

At room temperature, white prepupae were identified and marked within plastic food vials. 15-18 APF pupae were removed from the vial onto a piece of double-sided tape (Scotch brand, catalogue #665) applied to a microscope slide. Using fine forceps, the anterior pupal case was removed exposing the head and notum of all pupae applied to the tape, as previously described (O’Connor et al., 2022; Shannon et al., 2017). The tape was carefully removed from the microscope slide and inverted onto a pre-prepared cover glass (Corning 2980-246, 24 mm x 64 mm) (O’Connor et al., 2022). The pupae were carefully pressed down onto the cover glass by adhering the section of tape above the pupal head. Once the notum was visibly pressed onto the cover glass an oxygen permeable membrane (YSI, standard membrane kit, cat#1329882) was applied to prevent the pupae from drying out during imaging.

#### Pupal survival

Following imaging, pupae were kept mounted as described above and allowed to continue to develop and eclose for 3-4 days. Pupae that continued developing until they were able to crawl out of the partially dissected case were classified as ‘survived’ and their data acquired form these samples were used for analysis. If a pupae did not survive to eclosion, the associated datasets were not used in the study.

#### Live imaging

Images were collected using a Nikon Ti2 Eclipse with X-light V2 spinning disc (Nikon, Tokyo, Japan) with a 40X 1.3 NA oil-immersion objective or 60X 1.4 NA oil-immersion objective. Unless otherwise noted samples were imaged pre-wounding, immediately after wounding, every 2 min for 30 min, and then every 10 min for 6 h. Images were pre-processed in NIS-Elements using combinations of background subtraction, rolling ball correction, local contrast, and Denoise a.i. Assembly of figure panels was done using Affinity Designer and frames were centered on the entity in focus, compensating for frame shift due to wounding.

#### Laser ablation

A single pulse of a 3rd harmonic (355 nm) Q-switched Nd:YAG laser (5 ns pulse-width, Continuum Minilite II, Santa Clara, CA) was used for laser ablation. Laser pulse energies were kept to 1.9 μJ +/- 0.1 μJ, increased from our previous report (O’Connor et al., 2021b) to keep the wound size similar between old and new ablation rig.

#### Border breakdown and tangential vs radial assignment analysis

Individual border breakdown events were manually observed using FIJI by identifying syncytia late in the video and back-tracking to determine which borders broke down to form each syncytium. These borders were traced back to the first frame after wounding to develop the map on Fig 1G. For each border breakdown event, distance from the center of the wound and time after wounding were recorded in Microsoft Excel. These events were categorized as tangential or radial based on the orientation of the border-to-be-lost relative to a vector pointing outward from the center of the wound. Specifically, the line tool in FIJI was used to measure both the angle of the border with respect to the horizon, θ, and the angle of the line from the center of the wound to the center of the border with respect to the horizon, α. If |cos(*α* − *θ*)| ≥ cos 45°, then the border was classified as radial; otherwise, the border was classified as tangential.

#### Shrinking cell initiation / duration analysis

Shrinking cells were identified in live microscopy videos by beginning at the end of the video and playing backwards in FIJI. Backwards, a shrinking cell appears to bloom from the epithelial layer, characterized by the expansion of a bright puncta of p120ctn-RFP. Each cell that underwent this behavior was marked on a single frame of the video, then the distance from the center of the wound when shrinking started was denoted as well as the time that the shrinking started and completed. All cells that shrank were then manually traced back to the first frame after wounding to develop the map of shrinking cells (Fig 2E).

#### Approximating nuclei per syncytia

Nuclei and cell borders do not align in Z-projections of the pupal notum as the cells are non-prismatic. To estimate the number of nuclei per syncytium, the pre-wounding density of nuclei per unit area was determined for the circular region where syncytia form after wounding. Next, the apical area of the three largest syncytia/cells around a wound was measured at 0h, 1 h and 2 h post wounding. The area of each syncytia was multiplied by the nuclear density to yield the approximate number of nuclei per syncytia. The number of nuclei in each of the three syncytia was averaged to give a value for each of three samples in Figure 1B.

#### Measuring apical area of syncytia across varied wound sizes

The three largest syncytia were determined by eye in FIJI for six samples, three ablated at 1.9 µJ and three ablated at 3 µJ ablation. The apical area of syncytia was measured 3 h post wounding using the p120ctn-RFP signal. Initial wound size was calculated by measuring Myosin II marked leading edge when it became apparent 30-160 min post wounding.

#### Unfused cells at leading edge: count and percent analysis

Unfused cells at the leading edge were identified using FIJI by a lack of border breakdowns. Each unfused cell was manually observed over the duration of wound closure and the time at which it departed from the leading edge was noted. A count of unfused cells at the leading edge was created in Excel and formal graphs were generated using Prism 9. To measure percent of the leading edge comprised of unfused vs. syncytial cells, each unfused cell’s leading edge contact was measured in FIJI using the polygon line tool. The total circumference of the wound was measured using the polygon line tool and unfused cell measurements were subtracted from the total to infer the syncytial occupancy at the leading edge. Formal histograms were generated using Prism 9.

#### Analyzing GFP labeled cells

107 individually labeled GFP cells were analyzed across 5 wounds over 6.5 h. The position of the ablation was optimized to place as many individually labeled cells within 40-80 µm from the center of the wound as possible. After wounding it was possible to identify a mixing event by the decrease in intensity from the source cell with a corresponding increase in intensity of a previously unlabeled neighbor. Intensity differences made it possible to distinguish instances where two source cells were adjacent to each other but only one had a mixing event. However, large patches of source cells were not evaluated because inter-patch mixing was not distinguishable. To evaluate if border breakdowns were preceded by mixing, the 11 individually labeled cells that had border breakdowns were tracked back to the start of the video and confirmed to have a mixing event. There was never an instance where a labeled source cell had a border breakdown without a prior mixing event occurring.

#### Wound closure analysis

A pigmented scar forms at the site of laser ablation making identifying the exact moment a wound is closed difficult. Since the scar is approximately the same size in each sample, the time point at which the Myosin II signal disappeared below the scar was used as a proxy for closure.

#### Fusion analysis

A cell border was counted as a location of wound-induced fusion if it met three criteria: (1) adherens junctions along the cell border were lost; (2) the lost adherens junctions were not restored; and (3) the loss of adherens junctions was accompanied by a morphological change of the neighboring cells, for example, the moving apart of tri-cellular junctions previously connected by the lost cell border. Since border-breakdown fusions are observed most frequently within 1 hr after wounding, we counted all such fusions occurring within 1 hr 40 min of wounding.

When comparing the fusion frequency in *pnr* versus control domains, we used a landscape-oriented rectangle centered on the wound with its height defined by the inner radius of the zone of nuclear membrane damage visualized by the release of nls-mCherry from the nucleus, which is approximately the same as the zone of lysis (O’Connor et al., 2021a). Replacing the annular region with an extended rectangle excludes possible heterotypic fusions along the *pnr*-control border just above and below the wound. To calculate the frequency of fusion, in addition to the raw number of fusion borders, the total number of cell borders was also estimated by area-based scaling of the border count within a randomly selected 60-x-60-pixel (16.8-x-16.8-µm) region in the pre-wound image.

Since fusions can also appear as apical shrinking rather than border breakdown, we also quantified the number of apical-shrinking cell fusions in *pnr* versus control domains. Apical-shrinking cell fusions were defined as diploid cells that decreased in surface area over time and fully disappeared within a 5-hr movie. As with fusion analysis, cells near the *pnr*-control border were excluded. Apical shrinking cells were counted manually and their spatial locations marked using FIJI’s “Multi-point” tool.

#### Determining the effect of syncytia on wound closure

The distance between the leading edge and the wound center over time was used to calculate the speed of wound closure. The distance was calculated using the coordinates of the wound center and fixed points on the leading edge. To identify fixed positions on the leading edge that are unbiased and consistent over time, four reference lines, all crossing the center of the wound, were drawn using the red channel with only the nuc-mCherry signal: line 1 along the edge of the *pnr* domain labeled by nuc-mCherry, line 2 perpendicular to line 1, and line 3 and 4 are 30° from line 2. These references lines were then overlaid on the green channel with the E-cadherin signal.

To determine the speed of wound closure, the intersection of E-cadherin marked leading edge and reference lines 2-4 were used as the position of the leading edge. That is, at each time point, a total of 6 positions of the leading edge were recorded, among which 3 will be in the *pnr* domain and the other 3 in the ctrl domain. The average distance (d) between the leading edge and the center of the wound were calculated at each time point for both the *pnr* domain (d_pnr_) and the control domain (d_ctrl_). The difference between control and *pnr* distance (Δd = d_pnr_ – d_ctrl_) was calculated over time for both the control pupae with no genetic manipulation in the *pnr* domain (*pnr > +*) and experimental animals with *Atg1* knocked down in the *pnr* domain (*pnr > Atg1^RNAi^*). To determine whether Δd is statistically different between control and experimental pupae at each time point, two-way ANOVA (fit using full model) was used, comparing each cell mean with the other cell mean in that row.

To determine the distance syncytia covered during wound healing, syncytia were analyzed (1) at the leading edge when wound is half closed and (2) within the bow-tie region between reference line 3 and 4. ROIs were drawn along syncytia of interest using the “Freehand selections” tool of FIJI. In addition to syncytia location when wound is half closed, the starting position of cells that later fuse to form syncytia were also recorded in live-imaging videos, tracking the syncytia back in time. In addition to area, the coordinates of ROIs outlining syncytia at different time points were obtained using a self-written FIJI Macro. Within each ROI, the coordinate closest to the center of the wound was recorded using python, as the location of the syncytia. The distance a certain syncytium moved was calculated by subtracting the distance between the syncytium and the wound center when the wound was half closed from the distance between the syncytium and the wound center immediately after injury.

#### Calculating intercalation in live imaging

The change in the number of cells when the leading edge forms at 30 min (N_start_) to when the wound is closed (N_end)_ is equal to the number of intercalations plus the number of radial fusion events. Thus, (intercalations = ΔN – radial fusions). For the same three samples used in Fig 4D,G, we determined ΔN from N_start_ at 30 min and N_end_ when the wound had closed. Radial fusions were tallied by manually observing border breakdown events between the leading-edge cells and intercalations were calculated. Each radial fusion was counted as one prevented intercalation event.

#### Calculating cell shape index

Images of pupae labeled with Ecad-GFP were segmented into individual cells with Cellpose (Stringer et al., 2021); cells that were not readily segmented were excluded from subsequent analysis. To avoid perimeter inflation due to pixelation, the boundaries of segmented cells were smoothed by taking the midpoint between every other pair of boundary pixels. Smoothing and calculation of perimeters, areas and cell shape indices (*P*⁄√*A*) were performed in Mathematica (Wolfram Research Inc., Champaign, IL).

#### Vertex models with cell fusion

Simulations of wound healing were run using vertex models implemented in Mathematica (Wolfram Research Inc., Champaign, IL) following the methods of Tetley et al. (2019)and using parameters that match experimental observations from the *Drosophila* pupal notum as detailed in Han et al. (2024) with the following exceptions: the mean line tension on each edge (Λ) was specified in terms of a target shape parameter (*ρ*_0_) such that **Λ** = −2*ρ*_0_Γ√*A*_0_, where the contractility (Γ) and target cell area (*A*_0_) were taken from Han et al. (2024) and *ρ*_0_ = 3.745 was chosen so that the actual cell shape index matched experimental pre-wound measurements; and the line tension variability (σ_m_) was decreased from 1.0 x 10^-2^ to 9.3 x 10^-4^ so that the intercalation rate in unwounded cell sheets matched experimental observations. Pre-wound configurations were generated by performing a Voronoi tessellation of a circular area with 320 randomly distributed seed points and then equilibrating the cell configuration without stochastic fluctuations until it reached a steady state. Wounds were introduced by removing all cell borders within a circular patch ∼8 cells across and removing the wound’s contributions to the energy Hamiltonian. To handle wound closure and fusion, the following additions were made to the model: (1) purse-string tension on the leading edge of the wound was incorporated as in Tetley et al. (2019) with a normalized strength of 0.115; (2) traction/crawling forces of cells on the leading edge of the wound were incorporated as a uniform tensile stress acting inwardly on the edges and vertices of the wound area (similar to the uniform stress applied to the outer edge of the patch to represent tension from surrounding cells); this inward stress had a normalized strength of Σ*_in_*⁄*KA*_0_ = 0.13, where *K* is the area elasticity modulus; (3) the cell-cell interfaces at which fusion would occur were selected stochastically with probabilities that depended on time, distance from the wound, and the radial or tangential orientation of the interface – all chosen to match experimental observations reported here; and (4) the fusion of two cells was handled by eliminating the interface between those cells and setting the target area of the fused cell to the sum of the target areas of its two progenitors. All cells, fused or not, had the same contractility and target shape parameter.

#### Kymograph and plot profile analysis

Actin-GFP intensity was analyzed using the kymograph tool in FIJI after drawing a 11-pixel line through the middle of the syncytia. Profile plot values were exported from NIS elements to Excel and graphs were generated using Prism.

Key Resources Table

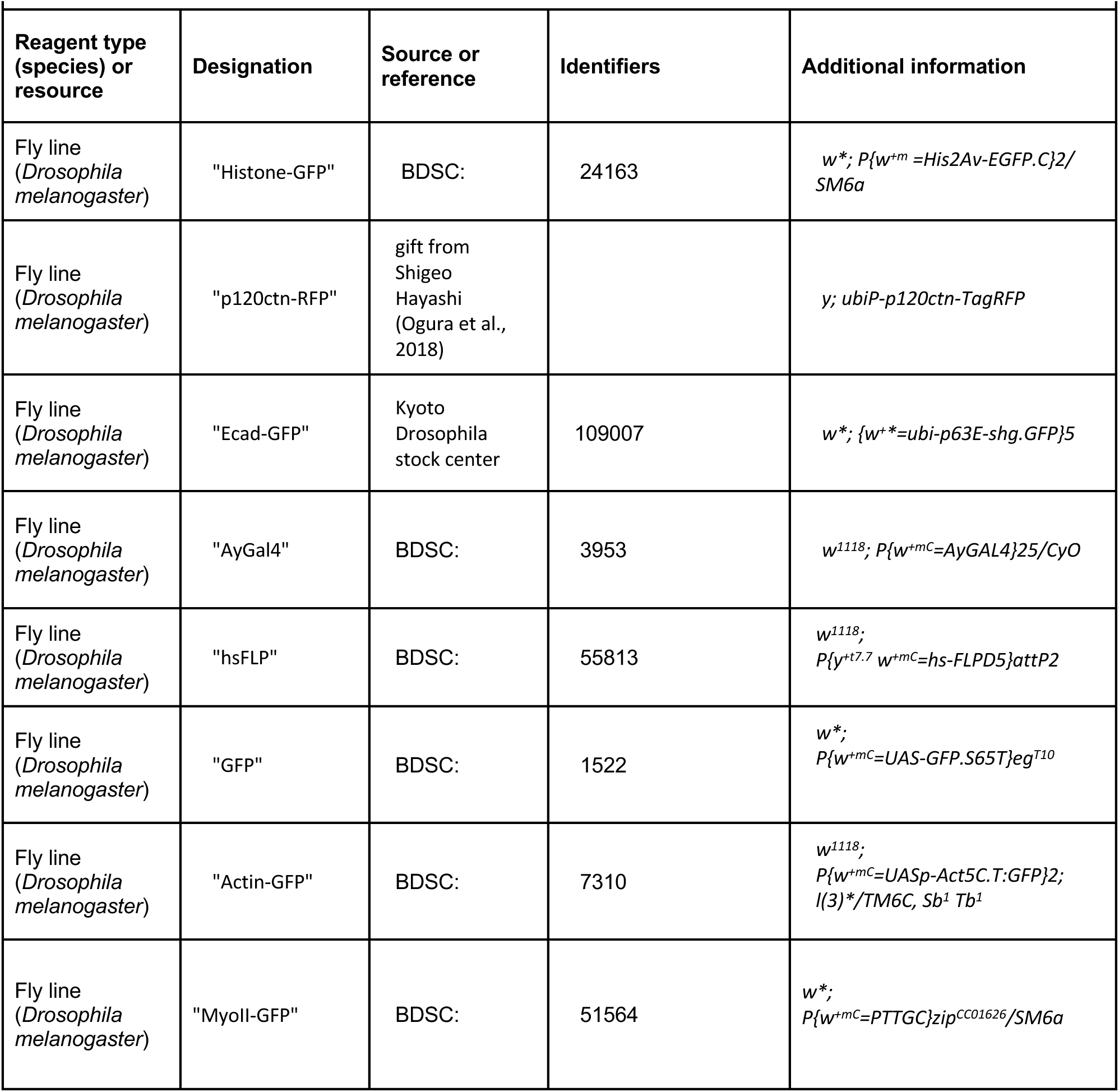
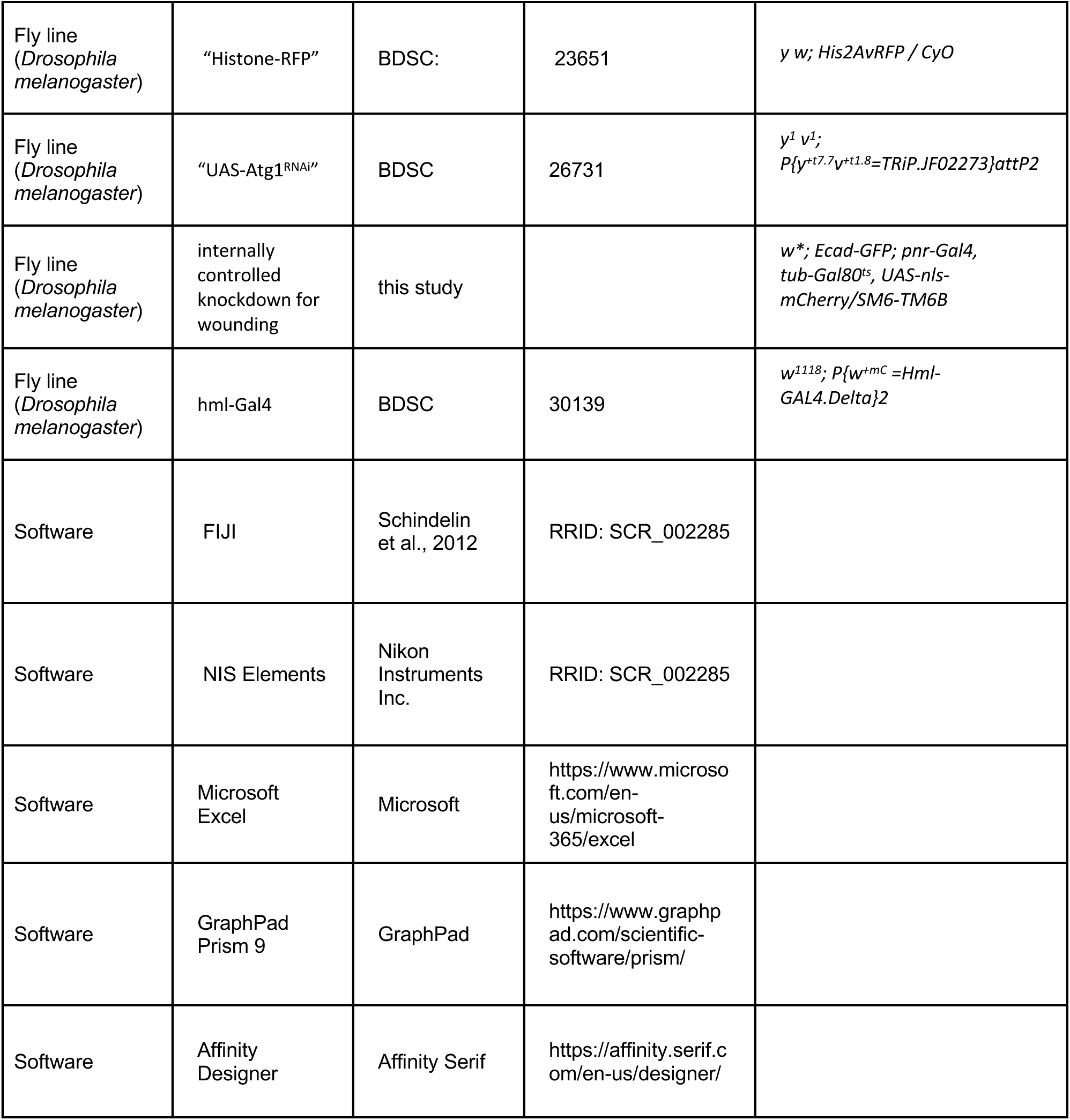

---

**Figure 1-figure supplement 1:**
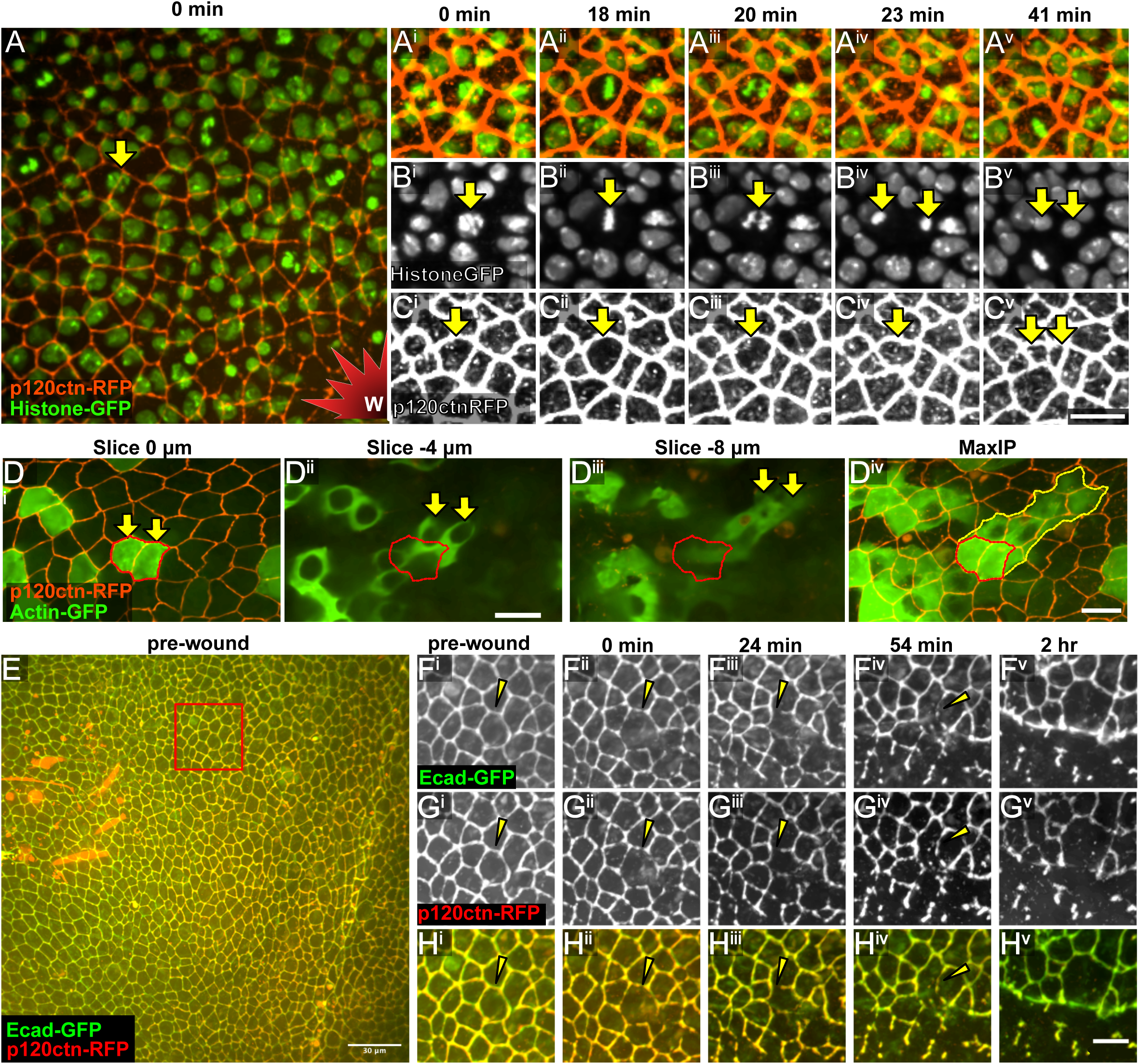
Characteristics of the pupal notum epithelium. **A-C)** An example of mitosis occurring ∼80 µm from the wound. Scale bar for A-C shown in C^v^, 10 µm. **D)** Z-slices at different depths of GFP-labeled cells reveal that the apical area (red outline) does not reflect the position of cells at basal slices (yellow arrows). The nuclear shadow in panel D^ii^ demonstrates the location of the nucleus. Scale bar, 10 µm. **E-H)** Ecad-GFP and p120ctn-RFP colocalize and behave similarly during border breakdown. Arrowheads in F-H points to borders breaking down in first hour after wounding. Scale bar for F-H shown in H^v^, 10 µm.

**Figure 3-figure supplement 1:**
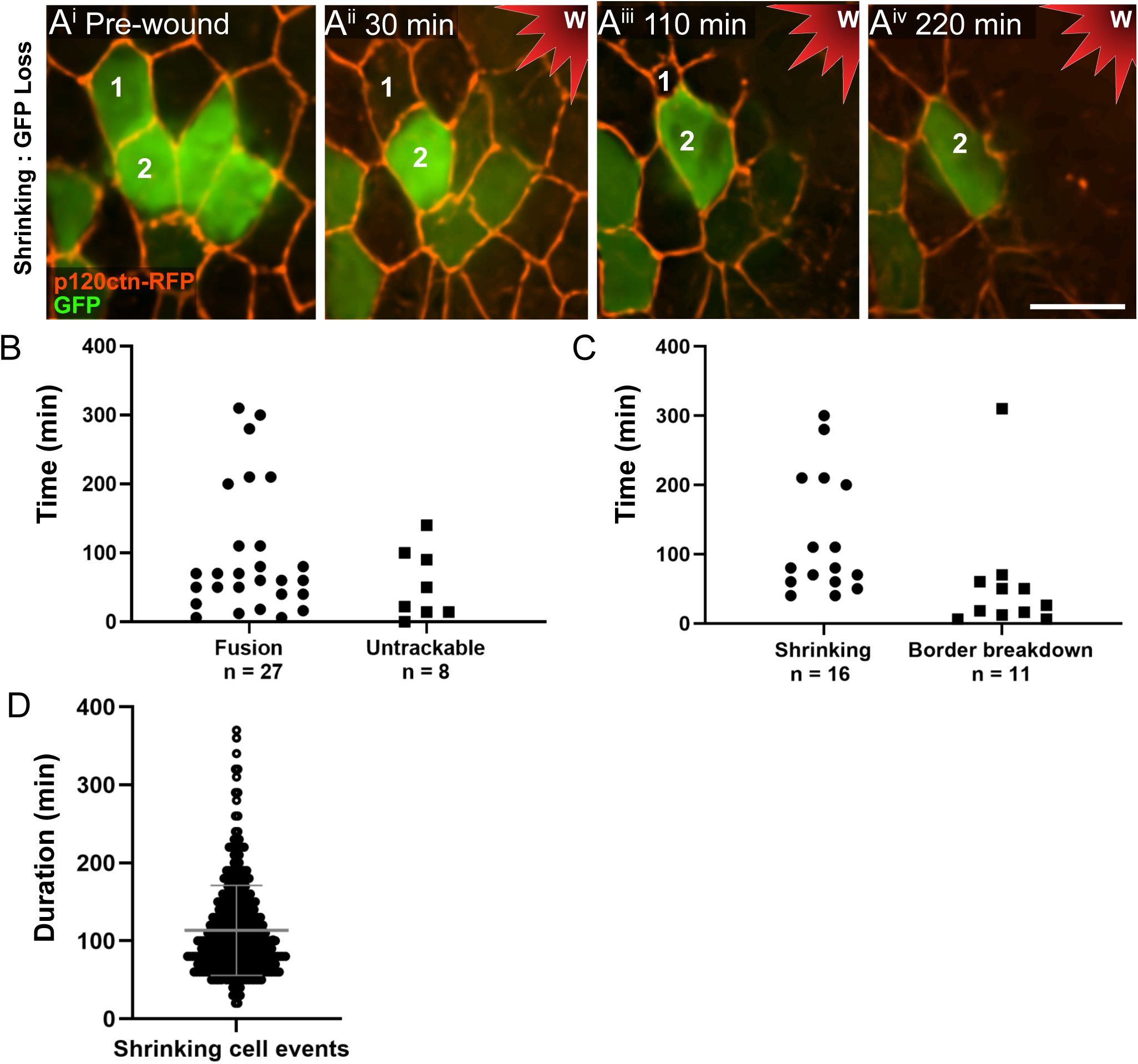
Temporal analysis of fusion events. **A)** An example of an untrackable cell is shown. After wounding, cell 1 loses cytoplasmic GFP, but there is no obvious recipient cell. Cell 1 then shrinks. Scale bar = 10 µm. **B)** Cells individually labeled with GFP reveal the timing and frequency of fusion or GFP loss (untrackable). **C)** Fusion cells from panel B were divided into two types of fusion events, shrinking cells and border breakdowns, to compare the temporal onset of each type of fusion. **D)** Cell shrinking was a lengthy and variable process, lasting up to hundreds of minutes. 274 shrinking cell events from three wounds were identified by p120ctn-RFP, mean and SD indicated.

**Figure 6-figure supplement 1:**
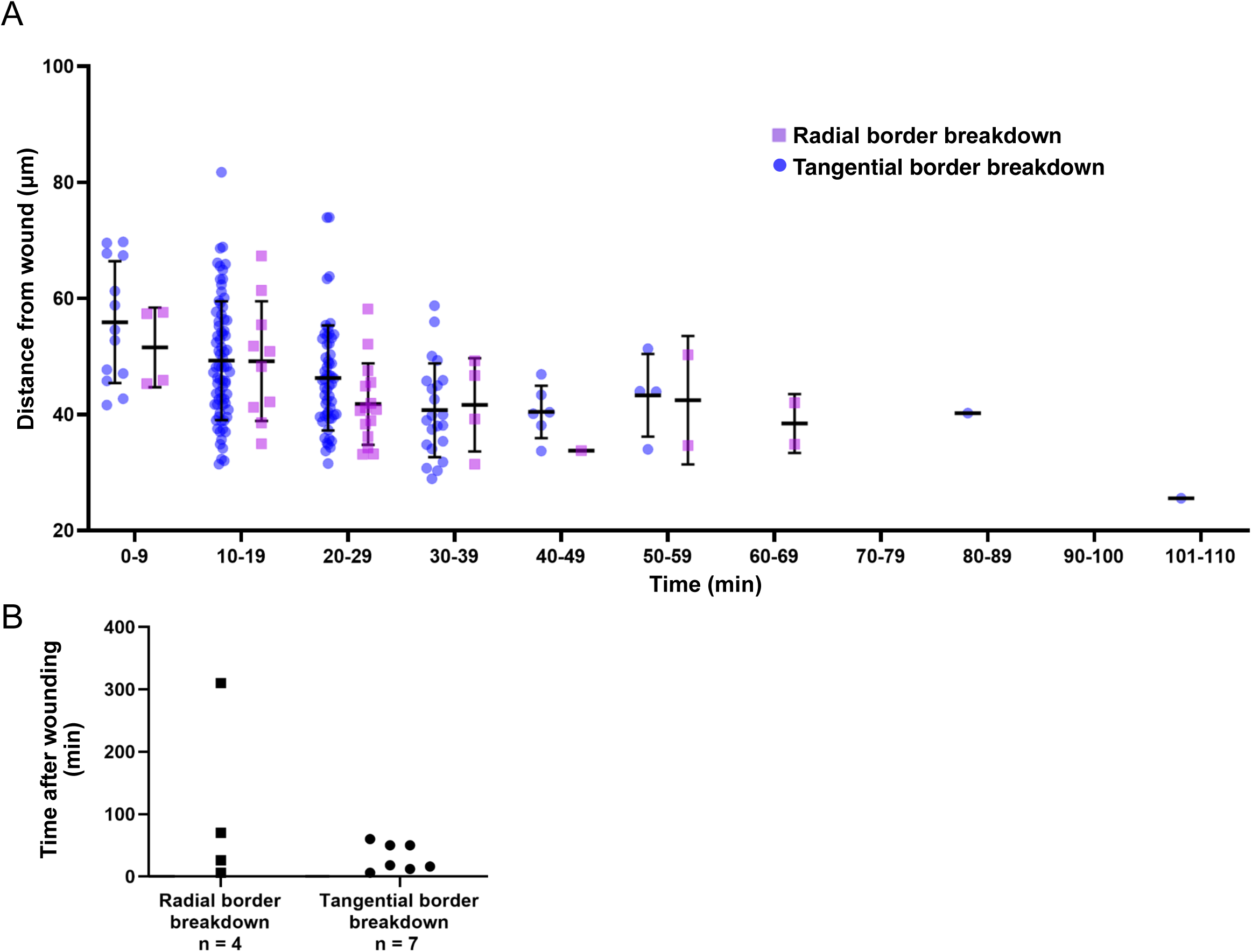
Comparisons of tangential and radial border breakdowns. A) The distance of tangential and radial border breakdown events from the wound, compared over time binned into 10 min intervals. The data is the same as in Figure 6B, from four wounds, with a total of 235 border breakdowns: 39 radial and 196 tangential. Border breakdown events were identified by p120ctn. **B)** The timing of tangential vs radial border breakdown fusion events observed in single GFP-labeled cells.

